# Global mapping of the energetic and allosteric landscapes of protein binding domains

**DOI:** 10.1101/2021.09.14.460249

**Authors:** Andre J. Faure, Júlia Domingo, Jörn M. Schmiedel, Cristina Hidalgo-Carcedo, Guillaume Diss, Ben Lehner

**Affiliations:** Center for Genomic Regulation (CRG), The Barcelona Institute of Science and Technology, Doctor Aiguader 88, 08003 Barcelona, Spain; Universitat Pompeu Fabra (UPF), Barcelona, Spain; Institució Catalana de Recerca i Estudis Avançats (ICREA), Passeig Lluís Companys 23, 08010 Barcelona, Spain

## Abstract

Allosteric communication between distant sites in proteins is central to nearly all biological regulation but still poorly characterised for most proteins, limiting conceptual understanding, biological engineering and allosteric drug development. Typically only a few allosteric sites are known in model proteins, but theoretical, evolutionary and some experimental studies suggest they may be much more widely distributed. An important reason why allostery remains poorly characterised is the lack of methods to systematically quantify long-range communication in diverse proteins. Here we address this shortcoming by developing a method that uses deep mutational scanning to comprehensively map the allosteric landscapes of protein interaction domains. The key concept of the approach is the use of ‘multidimensional mutagenesis’: mutational effects are quantified for multiple molecular phenotypes—here binding and protein abundance—and in multiple genetic backgrounds. This is an efficient experimental design that allows the underlying causal biophysical effects of mutations to be accurately inferred *en masse* by fitting thermodynamic models using neural networks. We apply the approach to two of the most common human protein interaction domains, an SH3 domain and a PDZ domain, to produce the first global atlases of allosteric mutations for any proteins. Allosteric mutations are widely dispersed with extensive long-range tuning of binding affinity and a large mutational target space of network-altering ‘edgetic’ variants. Mutations are more likely to be allosteric closer to binding interfaces, at Glycines in secondary structure elements and at particular sites including a chain of residues connecting to an opposite surface in the PDZ domain. This general approach of quantifying mutational effects for multiple molecular phenotypes and in multiple genetic backgrounds should allow the energetic and allosteric landscapes of many proteins to be rapidly and comprehensively mapped.

## Introduction

Proteins with important functions are usually ‘switchable’, with their activities modulated by the binding of other molecules, covalent modifications or mutations outside of their active sites. This transmission of information spatially from one site to another in a protein is termed allostery, which Monod famously referred to as ‘the second secret of life’^1–3^. Allosteric regulation is central to nearly all of biology, including signal transduction, transcriptional regulation, and metabolic control. Allosteric control can occur with or without large conformational changes and in proteins ranging from multisubunit protein complexes to individual protein domains. Many disease-causing mutations, including numerous cancer driver mutations, are pathological because of their allosteric effects^4,5^. Conversely, many of the most effective therapeutic agents do not directly inhibit the active sites of proteins but modify their activities by binding to allosteric sites. Amongst other benefits, allosteric drugs often have higher specificity than orthosteric drugs that bind active sites conserved in protein families^6,7^.

Allosteric sites are difficult to predict, even for highly studied proteins with known active and inactive states^8^. Individual proteins may contain a limited number of allosteric sites, which would be consistent with their physiological regulation by a limited number of ligands and modifications. Alternatively, as has been suggested by theoretical work, allostery might be quite widely distributed throughout protein domains^7–11^. This distinction between ‘sparse’ and ‘abundant’ allosteric sites has important implications: abundant allosteric sites would both facilitate the evolution of allosteric control^12^ and increase the likelihood of identifying therapeutic molecules that can bind a target protein and regulate its activity^13^. Most known allosteric sites are involved in physiological regulation but ‘orphan’ or ‘serendipitous’ sites without any understood physiological role have been identified for some proteins. An example is a cryptic allosteric pocket in the oncoprotein KRAS targeted by the first clinically approved RAS inhibitor^14^. Moreover, domain-insertion and mutagenesis also suggest quite extensive long-range communication in protein interaction domains^15^, enzymes^16–20^, transcription factors^21–23^ and receptors^24,25^.

Physical interactions between proteins are critical to most biological processes and, with >50,000 physical interactions between human proteins now identified and a total estimated interactome of >500,000 edges^6^, protein-protein interactions (PPIs) represent a vast and largely untapped therapeutic target space^6^. However, protein interaction interfaces are considered difficult drug targets^26^. When protein interactions can be allosterically regulated, this provides an alternative strategy for their therapeutic inhibition. However, allosteric sites are not known for most PPIs, a comprehensive map of allosteric sites has not been produced for any protein interaction domain, and generic methods to identify allosteric sites regulating PPIs do not exist.

Global maps of allosteric communication could be generated for protein binding domains if the effects of all mutations on binding affinity could be quantified: any mutation altering binding affinity but not directly contacting a ligand must be having an allosteric effect. However, changes in affinity cannot be inferred simply by quantifying changes in binding to an interaction partner; even in the simplest genotype-to-phenotype (energy) landscapes, ‘biophysical ambiguities’^27^ exist, meaning that changes in a molecular phenotype (e.g. binding to an interaction partner) can be caused by many different changes in the underlying biophysical properties (e.g. folding or binding affinity)^27,28^. To quantify the effects of mutations on binding affinity and so globally map allosteric communication, these ambiguities must be resolved.

Here we present an approach to achieve this for PPIs, allowing us to globally map the energetic and allosteric landscapes of protein interaction domains. The approach takes advantage of the massively parallel nature of deep mutational scanning (DMS) to quantify the phenotypic effects of thousands of perturbations^29^. We use an experimentally efficient strategy that we refer to as ‘multidimensional mutagenesis’ whereby the effects of mutations are quantified for multiple molecular phenotypes and in multiple genetic backgrounds. This method resolves ambiguities where a number of causal biophysical changes could account for an observed mutational effect^27,28^ and allows the inference of the *in vivo* biophysical effects of mutations. We harness the flexibility of neural networks to fit thermodynamic models to these experimental measurements (output) as a function of the corresponding mutagenised sequences (input), thereby accurately inferring the underlying causal changes in free energy. Applied to two protein domains, the method provides near complete views of their free energy landscapes and the first global maps of allosteric mutations.

## Results

### *ddPCA* quantifies the effects of mutations on protein abundance and binding

The binding of a protein to an interaction partner depends on both its affinity and the concentration of the active folded state. Allosteric sites altering binding affinity can not, therefore, be identified simply by quantifying how a perturbation changes the amount of protein bound to an interaction partner^30^—such an approach generates false positives where changes in binding are caused by changes in concentration and false negatives where changes in affinity are masked by changes in abundance. We therefore developed a strategy that uses two separate selection assays based on protein fragment complementation (PCA) to quantify the effects of mutations on both the abundance of a protein and its binding to an interaction partner (Figure 1a). As perturbations to probe the potential for allosteric regulation we use mutations which are a convenient method to introduce diverse changes in chemistry at all sites in a protein^31,32^. In the first assay, *bindingPCA*, the binding between two proteins is quantified by fusing them to different fragments of a reporter enzyme, dihydrofolate reductase (DHFR). Interaction between the proteins brings the DHFR fragments in close proximity allowing them to form a functional enzyme whose activity as measured by cellular growth in selective conditions is proportional to the concentration of the protein complex^33,34^. In the second assay, *abundancePCA*, only one protein is expressed and fused to a DHFR fragment with the other DHFR fragment highly expressed. Functional DHFR is now reconstituted by random encounters and growth is proportional to the concentration of the first protein over >3 orders of magnitude, as validated by applying the assay to >2000 yeast proteins^35^. We refer to the combination of these two assays as *DoubleDeepPCA* (*ddPCA*), a high-throughput method that quantifies the effects of mutations on both the abundance of a protein and its binding to one or more interaction partners.

**Figure 1.**
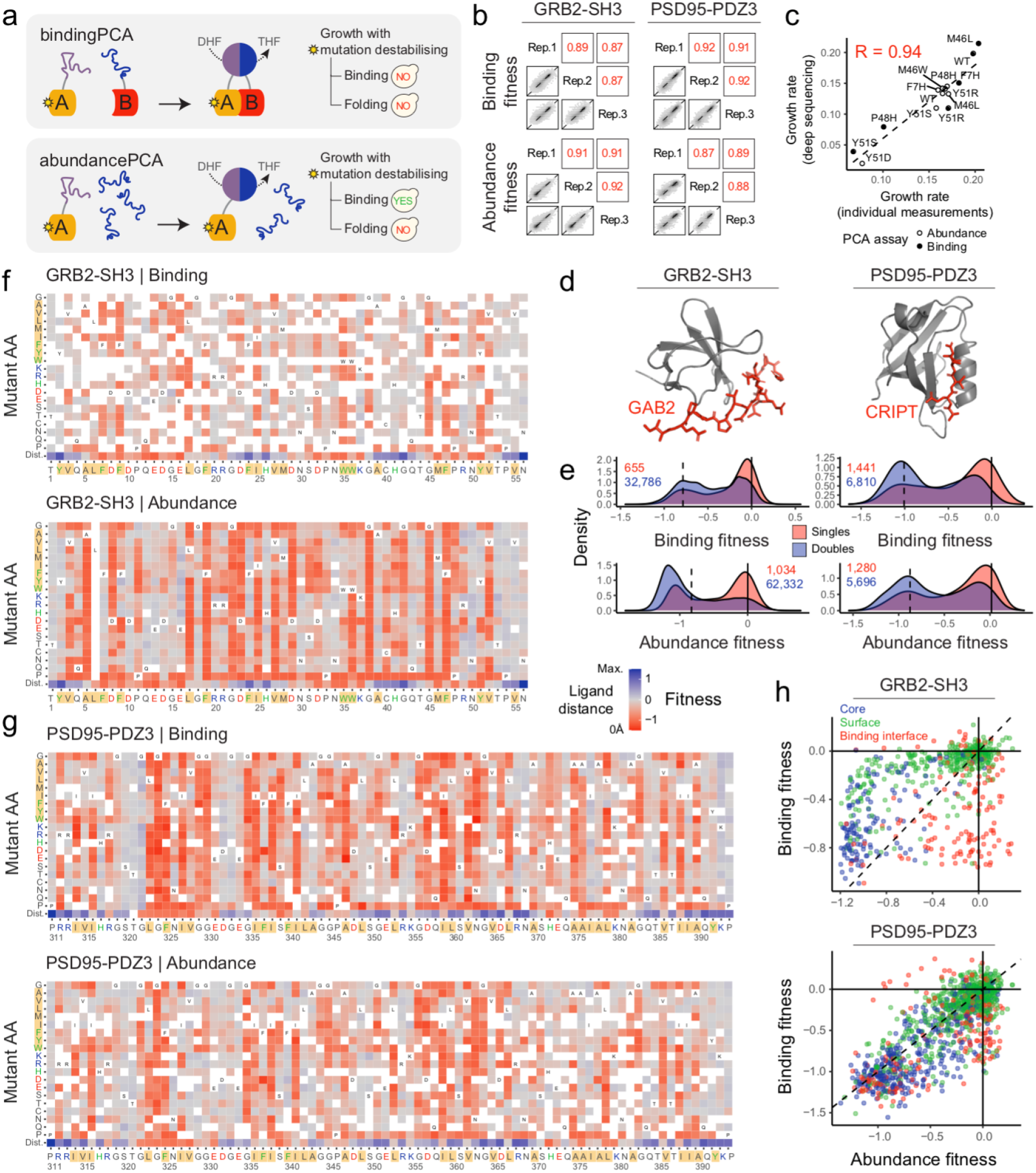
*ddPCA* quantifies the effects of mutations on protein abundance and binding. **a.** *ddPCA* uses two protein fragment complementation (PCA) selection assays to quantify the effects of mutations on the abundance (*abundancePCA*) of a protein of interest A and its binding to an interaction partner B (*bindingPCA*) based on growth in the presence of methotrexate. **b.** Reproducibility of fitness estimates from *ddPCA* for single and double AA substitutions between all biological replicates. Pearson correlations are indicated in red. **c.** Comparison of individually measured growth rates (slope of a linear fit of the log_10_(OD_600nm_) against time during the exponential phase) of GRB2-SH3 variants to growth rates inferred from deep sequencing data (see Methods). The dashed line corresponds to the linear regression model. R=Pearson correlation. **d.**3D structures of the SH3 domain of GRB2 bound to the ligand peptide of GAB2 (PDB entry 2VWF) and the third PDZ domain of PSD95 bound to the ligand peptide of CRIPT (PDB entry 1BE9). **e.** Fitness density distributions. Total variant counts for singles (red) and doubles (blue) are indicated (purple indicates overlapping density distributions). Vertical dashed lines indicate the median fitness of STOP codon mutations in the central 50% of the coding sequence. **f-g.** Heatmaps of the fitness effects of single AA substitutions for GRB2-SH3 (**f**) and PSD95-PDZ3 (**g**) corresponding to the *bindingPCA* (upper panel) and *abundancePCA* (lower panel) assays. The final row in each heatmap indicates the minimal distance between domain and ligand side chain heavy atoms (or alpha carbon atoms in the case of glycine). Amino acid labels are coloured in red or blue for positive or negatively charged residues respectively, green for aromatic amino acids and highlighted in yellow for hydrophobic residues. Fitness values more extreme than ±1.5 were set to this limit. **h.** Scatterplots comparing abundance and binding fitness of single AA substitutions. Variants are coloured by the corresponding residue position in the domain structure: core (relative solvent accessible surface area, RSASA<0.25), surface (RSASA≥0.25) or ligand binding interface (minimal side chain heavy atom distance<5Å).

We applied *ddPCA* to examples of two of the most common protein interaction domains encoded in the human genome: the C-terminal SH3 domain of the human growth factor receptor-bound protein 2 (GRB2), which binds a proline-rich linear peptide of GRB2 associated-binding protein 2 (GAB2)^36^, and the third PDZ domain from the adaptor protein PSD95/DLG4, which binds to the C-terminus of the protein CRIPT^37^.

There are two key principles of the *ddPCA* approach, which we refer to as ‘multidimensional mutagenesis’. First, the effects of mutations on two molecular phenotypes—binding and abundance—are quantified, and second, mutational effects are quantified starting from multiple genetic backgrounds. Both of these strategies are important for correctly inferring (disentangling) the underlying causal free energy changes from the measured mutational effects: many different free energy changes can generate the same change in phenotype^27^ and quantifying how mutations interact in double mutants^27,28,33^, as well as their effects on two different molecular traits, serves to resolve these biophysical ambiguities (Figure 2c). Moreover, the relationships between the free energies and folding and binding phenotypes/measurements are nonlinear and plateau at high and low energies^38^ (Figure 2f); quantifying the effects of mutations from different starting genotypes therefore serves to expand the effective dynamic range of individual measured mutational effects.

**Figure 2.**
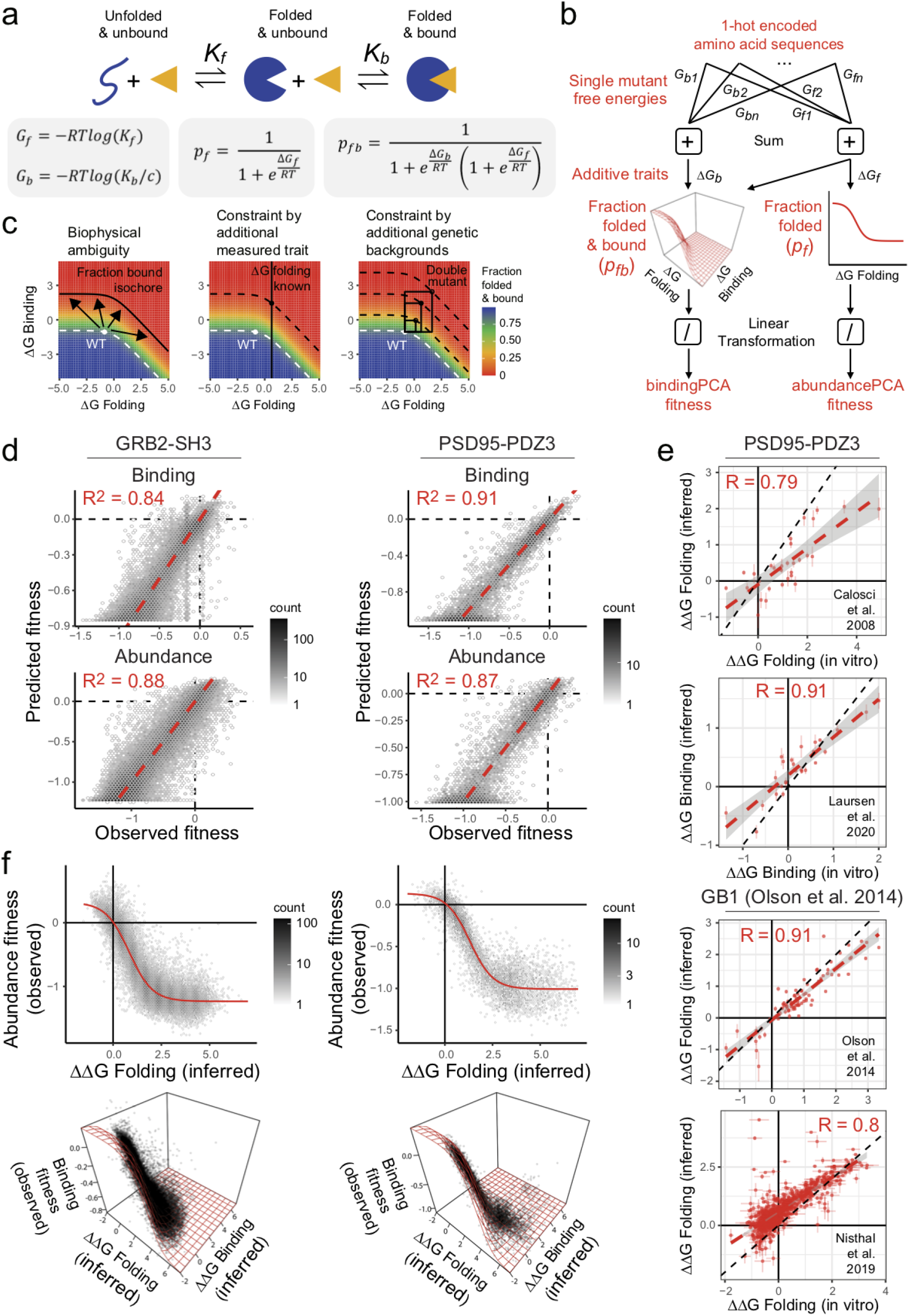
From molecular phenotypes to free energy changes. **a.** Three-state equilibrium and corresponding thermodynamic model describing relationships between binding and folding free energies and the fraction of bound protein complex or folded protein (see Methods). **b.** Cartoon depicting the neural network architecture used to fit the thermodynamic model to the *ddPCA* data (target/output data; bottom), thereby inferring the causal changes in free energy of folding and binding associated with single AA substitutions (input data; top). Also see Figure S2. **c.** Combinations of ΔG of binding and folding and the resulting fraction of bound protein complex (colour scale) illustrate how biophysical ambiguities (when different free energy change combinations result in the same phenotype; top panel) can be resolved by measuring more than one phenotype (middle panel) or by quantifying the effects of mutations in multiple starting genetic backgrounds (e.g. double AA substitutions; bottom panel). Curved lines indicate isochores i.e. distinct combinations of binding/folding free energy changes that give rise to the same molecular phenotype (bound fraction). **d.** Performance of models fit to *ddPCA* data comparing predicted and observed fitness for both measured phenotypes and protein domains. R^2^=proportion variance explained). **e.** Comparisons of the confident model-inferred free energy changes to previously reported *in vitro* measurements. R=Pearson correlation. Error bars indicate 95% confidence intervals. **f.** Non-linear relationships (global epistasis) between observed *abundancePCA* fitness and changes in free energy of folding (top panels) or *bindingPCA* fitness and both free energies of binding and folding (bottom panels). Thermodynamic model fit shown in red. Free energy changes outside the interval [−2,7] are not shown.

We generated mutagenesis libraries of the GRB2-SH3 and PSD95-PDZ3 domains containing both single and double amino acid (AA) substitutions (Figure S1a) and quantified their effects on binding to GAB2 and CRIPT, respectively, using *bindingPCA,* and on the concentration of the free domains using *abundancePCA*. All experiments were performed in biological triplicate, with deep sequencing used to quantify relative changes in binding and abundance in pooled selection assays (Figure S1b,c). We calculated abundance and binding fitness scores and associated errors using DiMSum^39^. Binding and abundance fitness scores were highly reproducible between replicates (Figure 1b, Pearson’s R = 0.87-0.92). Mutational effects also agreed very well with individual growth measurements (Pearson’s R = 0.94, p-value = 5.1e-07, Figure 1c).

The distributions of mutational effects corresponding to both binding and abundance are bimodal for both domains, with, for example, 28% of single AA substitutions strongly affecting binding of the PDZ domain and 46% having nearly neutral or mild effects (*bindingPCA* fitness within the lower peak < −0.75 and upper peak > −0.25 respectively, Figure 1e). The mutational effect matrices for binding reveal that mutations with large effects on binding are distributed throughout both domains (Figure 1f-g). Similarly, the mutational effect matrices for abundance show that mutations throughout both domains also have large effects on protein concentration (Figure 1f-g). Indeed, plotting the changes in binding against the changes in abundance reveals that most mutations altering binding also alter the concentration of the isolated domains (Figure 1h), consistent with the expectation that changes in protein stability are a major cause of mutational effects on binding^40^.

### From molecular phenotypes to free energy changes

We used a neural network formulation of the Boltzmann free energy equation to fit thermodynamic models to the experimental data obtained using *ddPCA*, thereby inferring the underlying causal free energy changes from the effects of each single AA substitution on these two molecular phenotypes (Figure 2a,b, Figure S2.5). Protein binding can be most simply modelled as a three-state equilibrium with unfolded, folded and bound energetic states (Figure 2a). In this genotype-phenotype model, mutations alter the free energy of folding (ΔG_f_) and/or binding (ΔG_b_), and changes in free energy combine additively in double AA substitutions. The relationship between the fraction of folded or folded+bound protein and the respective measured phenotypes (*abundancePCA* and *bindingPCA* fitness) is assumed to be linear^28,33^ (Figure 2b; see Methods).

The three-state model provides an excellent quantitative fit to the data for both domains (Figure 2d, R^2^ = 0.84-0.91, see Figure S3 for similar comparisons shown separately for single and double AA substitutions, as well as for held out validation data not used for training), strongly supporting the assumption that most changes in the free energy of both folding and binding are additive in double AA substitutions^32,41^. Training models using data corresponding to only one molecular phenotype (binding, Figure S4a), or two molecular phenotypes but only fitness effects from single AA substitutions (Figure S4b), results in worse fits to the data (R^2^ = 0.05-0.92 and R^2^ = 0.77-0.83 respectively). The number of double AA substitutions for each single AA substitution varies across the four datasets, with the relatively few double AA substitutions in the PSD95-PDZ3 libraries (median=5 and 4 in the binding and abundance libraries respectively, Figure S1a) still sufficient to infer the underlying free energy changes. Downsampling double mutant data in both datasets illustrates how increasing the number of double mutants improves the model fit (Figure S5).

To evaluate the quality of the inferred free energy changes upon mutation, we compared them to *in vitro* measurements for the PDZ domain. We find excellent agreement between inferred free energies of folding relative to the wild-type (ΔΔG_f_) and those corresponding to single AA substitutions determined *in vitro* for PSD95-PDZ3 (F337W background)^42^ (Figure 2e, Pearson’s R = 0.79, n = 30, p-value = 2e-7). Binding free energy changes similarly agree with those measured by stopped-flow experiments for the PSD95-PDZ3:CRIPT interaction^43^ (Figure 2e, Pearson’s R = 0.91, n = 26, p-value = 8e-11; see Figure S6 for additional comparisons to smaller-scale *in vitro* validation datasets). Using only the binding or only single AA substitutions data results in worse agreement with the *in vitro* binding and folding free energy changes (Figure S4), as does reducing the number of double mutants used to fit the model (Figure S5). As a further validation of our method to infer free energy changes from molecular phenotypes, we fit the same three-state model to previously published *in vitro* mutagenesis data for the binding of nearly all single and double AA substitutions of protein G domain B1 to IgG-Fc^44^ (Figure S3a,d). Even in the absence of multiple measured phenotypes (binding only), with this depth of double mutant data we find excellent agreement between inferred free energy changes of folding and *in vitro* measurements^44,45^ (Figure 2e, Pearson’s r = 0.8-0.91, n=685 and 80, p-value<2.2e-16), similar to a previous analysis^28^. These comparisons further demonstrate the validity of our inferences and the general flexibility of the approach.

### Binding and folding free energy landscapes of the SH3 and PDZ domains

Free energy landscapes of mutational effects (Figure 3a,b) have important advantages over maps of phenotypic effects (Figure 1f,g), converting mutational effects to the underlying additive biophysical traits and allowing accurate genetic prediction when mutations are combined in pairs and larger combinations^27^. Furthermore, the free energy landscapes are more complete, as single mutant free energies can be inferred by their effects in different backgrounds (e.g. double AA substitutions) or their effects on a related phenotype (e.g. folding energies from binding phenotype) despite missing single mutant phenotypes.

**Figure 3.**
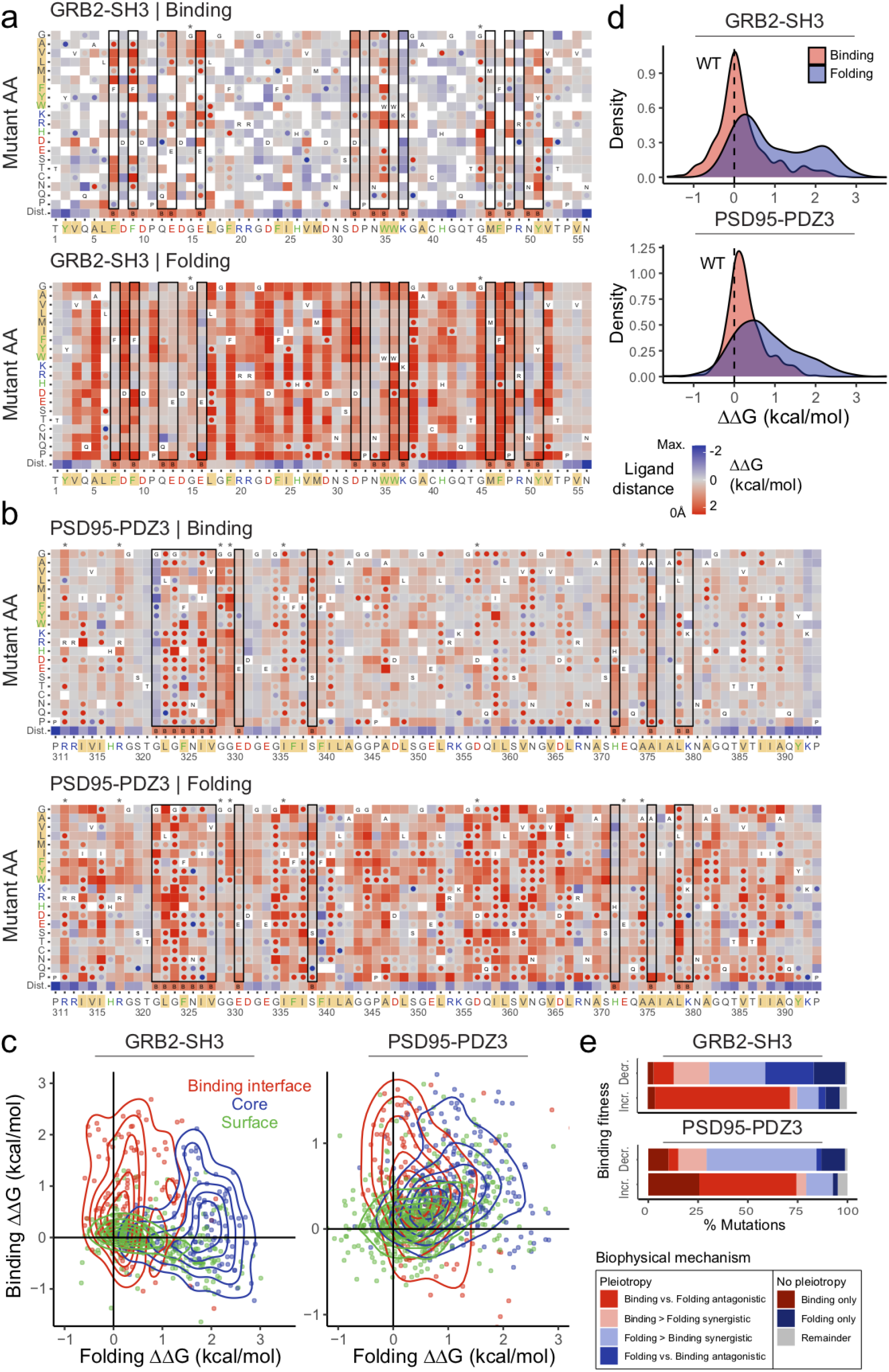
Binding and folding free energy landscapes of the SH3 and PDZ domains. **a-b.** Heatmaps showing inferred changes in free energies of binding and folding for GRB2-SH3 (**a**) and PSD95-PDZ3 (**b).** The final row in each heatmap indicates the ligand distance and amino acid labels are coloured according to the chemical structure and properties of their side-chains (see Figure 1f,g). Free energy changes of ligand-proximal residues (ligand distance<5Å) are boxed and asterisks indicate major allosteric positions. Lower confidence estimates are indicated with dots (95% confidence interval ≥1kcal/mol). Free energy changes more extreme than ±2.5kcal/mol were set to this limit. **c.** Scatterplots comparing confident binding and folding free energy changes of mutations in the core, surface and binding interface. Contours indicate estimates of 2D densities using 6 contour bins. Axis limits were adjusted to include the largest contour bin (more extreme data points are not shown). **d.** Distributions of confident binding (red) and folding (blue) free energy changes. X-axis limits were adjusted to match those in panel c. **e.** Percentage of mutations that significantly decrease (top) or increase (bottom) fitness in the binding assay (FDR=0.05) categorized by their biophysical mechanism. Pleiotropic mutations have significant changes in free energies of both folding and binding (FDR=0.05) and are classified as either synergistic or antagonistic depending on whether their effects are in the same or different direction respectively. See Figure S7 for the GB1 domain.

In general, mutations act asymmetrically on the two inferred biophysical traits, with mutations tending to have stronger effects on the free energy of folding than binding (Figure 3a,b,d, Figure S7a,b,d). This is not evident from comparisons at the phenotypic level (Figure 1f,g). Mutations that affect folding are also more numerous and more widely distributed throughout the domains, with many positions sensitive to perturbation. Effects on binding affinity on the other hand are comparatively less frequent and enriched in residues proximal to the ligand i.e. in the binding interface (Figure 3a,b, Figure S7a,b, black boxes). Thus, changes in protein binding (change in *bindingPCA* fitness) are predominantly driven by changes in protein stability, especially for mild and intermediate fitness effects (Figure S8a).

Directly comparing free energies stratified by the position of the mutated residues reveals that mutations in core residues (relative solvent accessible surface area, RSASA<0.25) have the largest effects on folding, whereas mutations in the binding interface (ligand distance<5Å) have the strongest effects on binding affinity (Figure 3c, Figure S7c). Surface residues (RSASA≥0.25) are more tolerant to mutations (Figure 3c, Figure S7c).

The bimodality of phenotypic effects (Figure 1e) is much reduced in distributions of relative free energy changes, particularly in the case of binding free energies, whose mode is centered on ΔΔG=0 i.e. no difference from wild-type (Figure 3d, Figure S7d). The distribution of folding free energies has a positive mode (ΔΔG>0) and a heavy right tail indicating that most single AA substitutions have destabilising effects and many substitutions are strongly destabilising (Figure 3d, Figure S7d).

### Extensive biophysical pleiotropy

Comprehensively quantifying the effects of mutations on both folding and binding provides the first opportunity to assess the extent to which mutations affect multiple biophysical properties i.e. biophysical pleiotropy^27^. Overall, more than two thirds of all mutations altering binding are biophysically pleiotropic (86% and 67% increasing binding and 80% and 77% of mutations decreasing it are biophysically pleiotropic in GRB2-SH3 and PSD95-PDZ, respectively). In both domains, mutations that disrupt binding most often show synergistic pleiotropy, reducing both binding and folding stability (Figure 3e, Figure S7e). In contrast, mutations that increase binding tend to display antagonistic pleiotropy with the effects on binding and folding free energies in opposing directions (Figure 3e, Figure S7e). The proportion of different kinds of pleiotropic mutations differs depending on the region of the domain (Figure S8b). For instance in GRB2-SH3 and PSD95-PDZ, compared to core or surface residues, binding interface positions harbour a higher proportion of mutations that disrupt binding despite antagonistic effects on the free energy of folding (see below), consistent with the hypothesis that residues of proteins involved in substrate binding or catalysis are not optimized for stability^46^.

This extensive biophysical pleiotropy further emphasises the importance of determining the biophysical effects of mutations. For both genetic prediction and protein engineering, the outcome when combining mutations is often only predictable if the causal biophysical effects can be measured or inferred: combining mutations with the same phenotypic outcomes but different biophysical causes often results in different phenotypic consequences^27^.

### Surface residues are suboptimal for protein stability

Overlaying the mean folding free energy changes per residue on the domain structures further illustrates that solvent exposed (surface or binding interface) residues tend to be less sensitive to mutations than those that constitute the buried core of the structure (Figure 4a,b, Figure S9a,b, two-sided Mann-Whitney U test p-value<2.2e-16), with per-residue mean folding energies anti-correlated with residue burial (RSASA; Figure 4c, Figure S9c).

**Figure 4.**
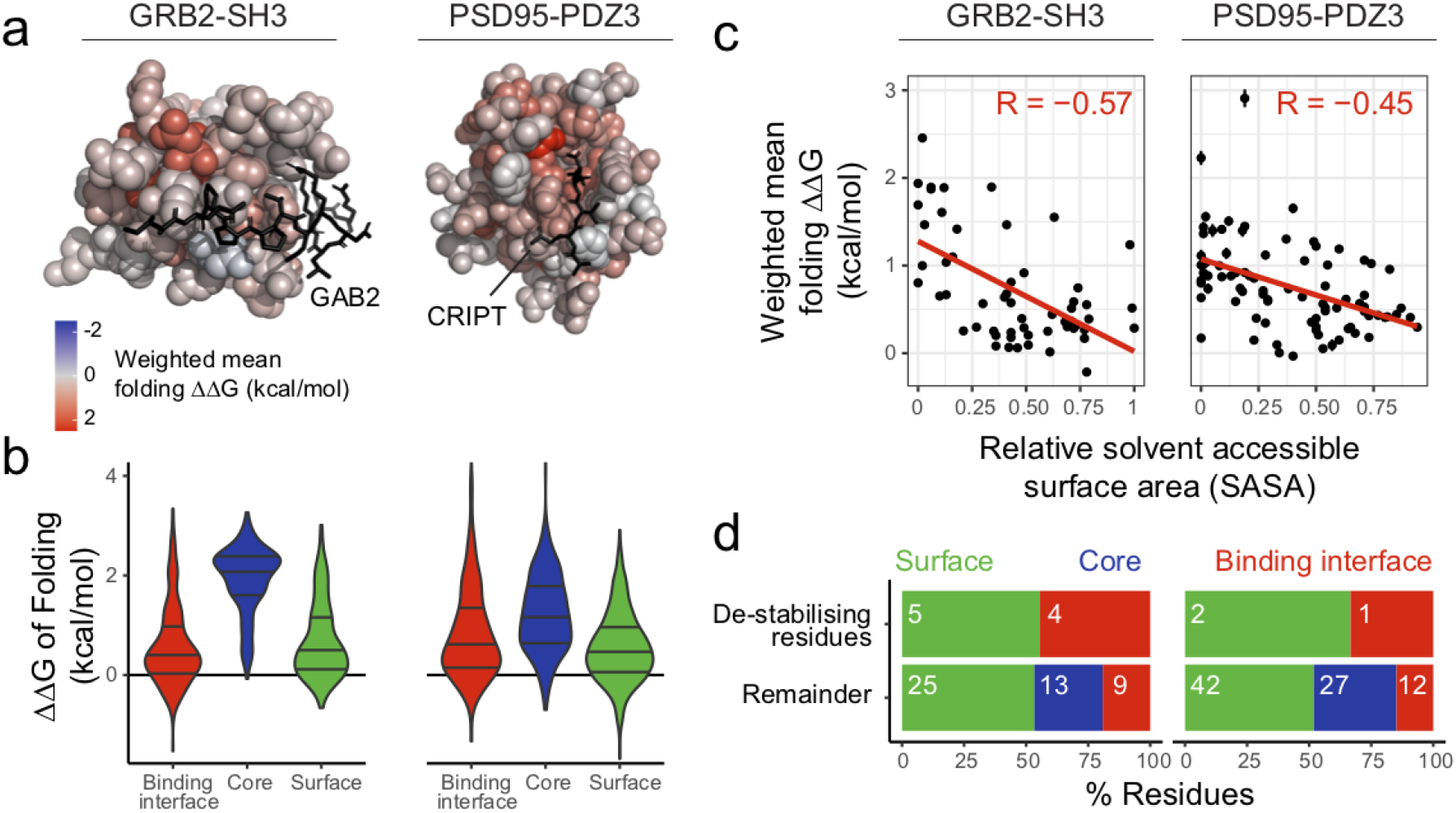
Mutational effects on protein stability. **a.** 3D structures of the GRB2-SH3 and PSD95-PDZ3 domains where residue atoms are colored by the position-wise average change in the free energy of folding. Ligands are shown as black sticks. **b.** Violin plots indicating distributions of confident changes in free energy of folding stratified by position in the structure (two-sided Mann-Whitney U test p-value<2.2e-16 comparing mutations in the core versus the remainder for both protein domains). **c.** Negative correlation between the position-wise average change in free energy of folding and the solvent exposure of the corresponding residue (RSASA). R = Pearson correlation. **d.** Percentage of core, surface or binding interface residues shown separately for de-stabilising residues (positions with ≥5 stabilizing mutations, folding ΔΔG<0, FDR=0.05) and the remainder. Inset numbers are total counts. See Figure S9 for the GB1 domain.

Although mutations overwhelmingly destabilise folding, a small subset of residues are enriched for mutations that increase stability (ΔΔG<0). We find nine residues in GRB2-SH3 and three residues in PSD95-PDZ3 where at least five distinct single AA substitutions are observed to decrease the free energy of folding compared to the wild-type, none of which are classified as core residues (Figure 4d). That AAs unfavourable for stability have been retained by evolution suggests selective constraints other than stability acting on these sites^46^. Consistent with this, 4/9 destabilising residues in GRB2-SH3 and 1/3 destabilising residues in PSD95-PDZ3 are in the binding interface (ligand distance<5Å). The remaining surface destabilising residues may facilitate binding to other partners, be sites of post-translational modification or be involved in other molecular functions. Indeed, all of these residues are either highly evolutionarily conserved (Figure S10a) or uncharacteristically hydrophobic compared to other surface residues (pooled two-sided Mann-Whitney U test p-value = 2e-2 for both protein domains), having hydrophobicity scores comparable to residues within the buried core of the domains (Figure S10b). This suggests that these residues are involved in additional molecular interactions. In GRB2-SH3, three surface destabilising residues, Y2, F24 and V55, form a hydrophobic cluster that is part of the dimer interface of full-length GRB2^47^ (Figure S11). Similarly, in PSD95-PDZ3 both of the surface destabilising residues, F340 and P394, are involved in extra-domain interactions with residues C-terminal to PDZ3 (Figure S11).

This illustrates how *abundancePCA* data can help identify functionally important surface sites, even when interaction partners and functions are unknown. Moreover, in general, identifying surface sites that are suboptimal for stability (but otherwise functionally neutral) has therapeutic implications, predicting where the binding of small molecules or biologics may help stabilise proteins carrying disease-causing destabilising mutations^48,49^.

### Binding interface identification

Overlaying the mean absolute binding free energy changes per residue on the domain structures shows a strong enrichment for the largest mutational effects in the binding interface (ligand distance<5Å, Figure 5a, Figure S12a). Indeed, the inferred binding free energy changes alone accurately predict binding interface residues (area under the ROC curve, AUC = 0.83-0.94, Figure S13a). Furthermore, the effects of individual mutations at key ligand-contacting residues are consistent with structural information at those positions (Figure S13b). This illustrates how quantifying the effects of mutations on binding and abundance (but neither of these phenotypes alone, Figure S13c) can identify binding interfaces, which may be useful for identifying the interfaces of the very large number of protein interactions without any structural information^50^.

**Figure 5.**
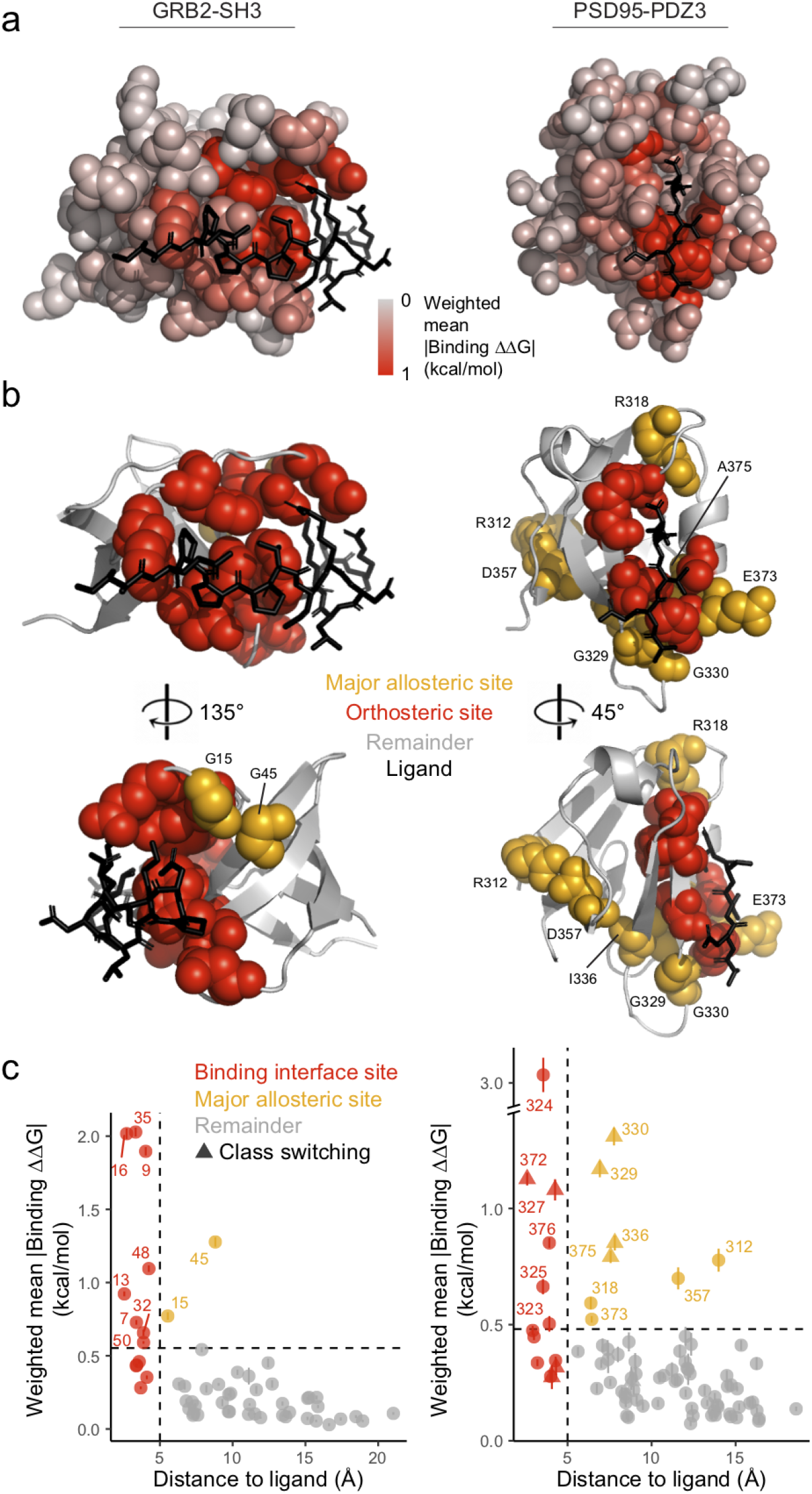
Major allosteric sites in protein binding domains. **a.** 3D structures of the GRB2-SH3 and PSD95-PDZ3 domains where residue atoms are colored by the position-wise average absolute change in the free energy of binding. Ligands are shown as black sticks. **b.** Domain structures with orthosteric sites (ligand distance<5Å) highlighted in red spheres and major allosteric site residues highlighted in orange spheres. **c.** Relationship between the position-wise average absolute change in free energy of binding and the distance to the ligand (minimal side chain heavy atom distance). Major allosteric sites (orange) are defined as non-binding interface residues with weighted average absolute change in free energy of binding higher than the average of binding interface residue mutations (red). Class-switching residues in PSD95-PDZ3 are those that favour a change in specificity for a T-2F ligand defined in McLaughlin et al. 2012^51^. See Figure S12 for the GB1 domain.

### Comprehensive maps of allosteric mutations

We next asked whether any residues outside the binding interface are also enriched for mutations that modulate binding affinity. We identified two sites in GRB2-SH3 (G15 and G45) and 8 sites in PSD95-PDZ3 (R312, R318, G329, G330, I336, D357, E373 and A375) with mean absolute change in binding free energy greater than the mean absolute change of all mutations in the binding interface (Figure 5b,c, Figure S14a,b). We refer to these ligand-distal residues at which many mutations have strong effects on binding affinity as major allosteric sites (see Figure S12b,c for the GB1 domain).

A previous study identified 9 residues within PDS95-PDZ3 at which AA substitutions have the capacity to switch PSD95-PDZ3 ligand binding class specificity—an observation that can only be explained by an underlying causal change in binding affinity^51^. 6/9 of these class-switching residues (two within and four outside the binding interface) are identified as major allosteric sites by our definition. Moreover, the remaining three class-switching residues (G322, V362 and L379) are enriched for mutations with strong (albeit low confidence) binding free energy changes compared to other residues not classified as major allosteric sites (two-sided Mann-Whitney U test p-value=1e-10, AUC=0.76). This identification of previously described specificity-determining residues as major allosteric sites further validates our approach.

The two major allosteric sites in GRB2-SH3 and 6/8 in PSD95-PDZ3 (R318, G329, G330, I336, E373, A375) are either in direct physical contact with binding interface residues (i.e. second shell sites) or immediately adjacent to them in the linear AA sequence (i.e. backbone-backbone contacts; see asterisks in Figure 3a,b). However, two sites in PSD95-PDZ3 (R312 and D357) are on the opposite surface of the domain, with distances of 11-14Å to the closest ligand residue. These sites form a near-contiguous “chain” of residues linking the N-terminal beta strand to the binding interface via a salt bridge formed by residues R312 and D357, where the latter is in close proximity to the second-shell class-switching residue I336 located in the adjacent beta sheet strand (minimum backbone atom distance=4.1Å). Thus, whilst the sites most enriched for mutations affecting binding affinity are mostly proximal to the binding interface, in the PDZ domain they also extend throughout the domain to the opposite surface.

### Allosteric mutations are distributed throughout protein cores and surfaces

Although these 10 residues are the positions most enriched for allosteric effects, mutations affecting binding affinity actually occur throughout both protein domains (Figure 3a,b). Defining allosteric mutations as those with effects at least as large as the mean absolute binding free energy change of mutations in the binding interface, we find a total of 55 allosteric mutations in 24 distinct residues in GRB2-SH3 (33 core, 22 surface) and 152 allosteric mutations in 49 residues in PSD95-PDZ3 (83 core, 69 surface). 40% (12/30) and 55% (24/44) of all surface residues have at least one allosteric mutation in GRB2-SH3 and PSD95-PDZ3 respectively (Figure 6a,b, Figure S15, Figure S16a,b). These results suggest that allosteric mutations are abundant in the core of proteins and also in solvent accessible regions. Moreover, similar to their sub-optimality for folding, surface sites are also often suboptimal for binding at a distal interface.

**Figure 6.**
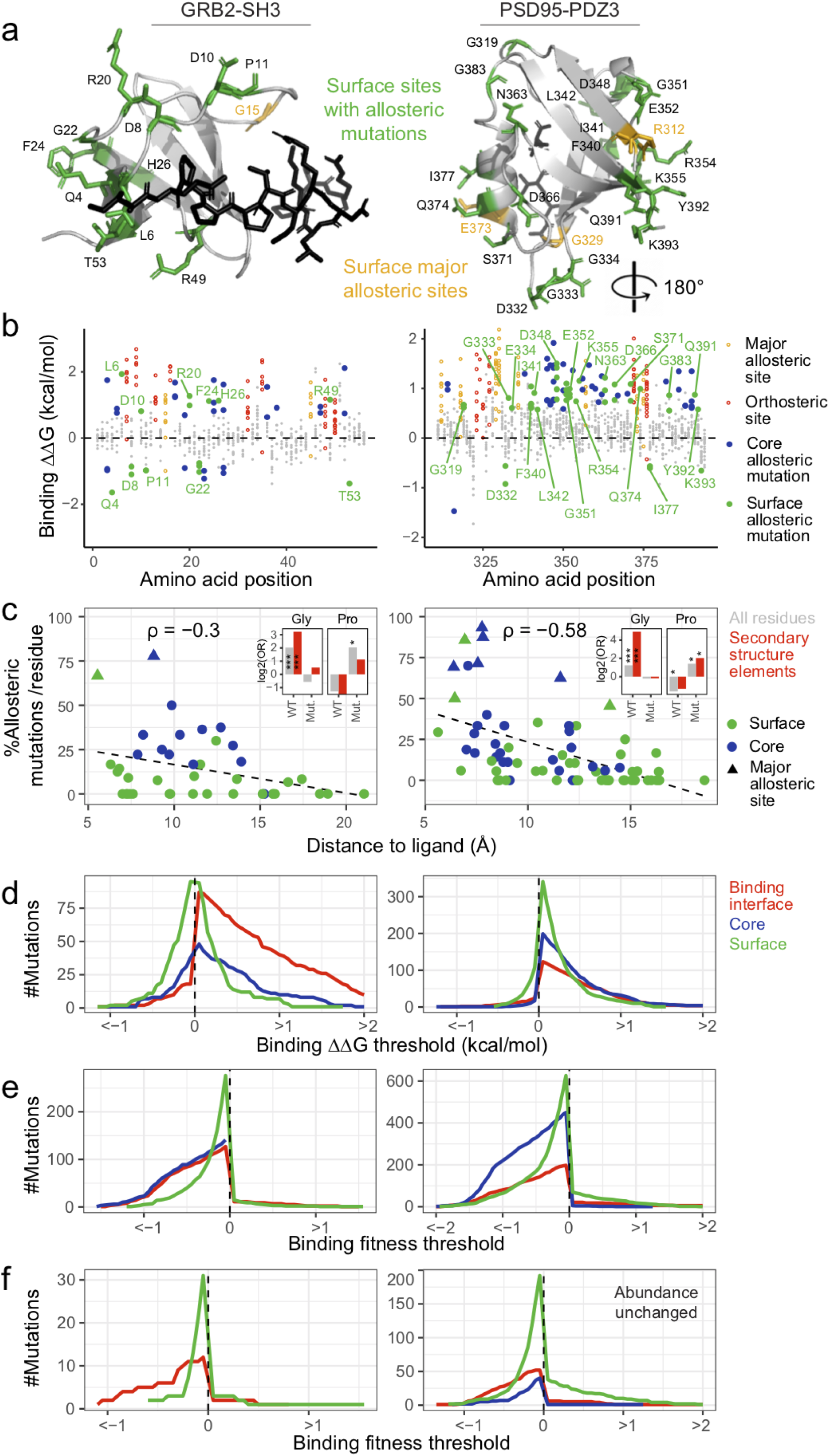
Protein surfaces are frequent sites of binding affinity modulation. **a**. Domain structures with surface allosteric sites and surface residues with allosteric mutations highlighted in orange and green respectively. **b**. Scatterplot showing the binding free energy changes of all mutations and coloured according to residue position: allosteric site (orange), orthosteric site (red), core allosteric mutation (blue), surface allosteric mutation (green). **c.** Percentage of allosteric mutations per residue versus ligand proximity, excluding sites within the binding interface (ligand distance<5Å). Points are colored according to residue position and major allosteric sites are indicated (see legend). ρ=Spearman rank correlation coefficient. Inset: enrichment of allosteric mutations at WT (or introducing mutant, Mut.) Glycines and Prolines in positions outside the binding interface or further restricted to those in secondary structure elements. The log2 odds ratio corresponding to a 2×2 contingency table is shown on the y-axis and the associated P-value from a two-sided Fisher’s Exact Test is indicated (*P-value<0.05, ***P-value<0.001). Also see Figure S17b,c. **d.** Total numbers of mutations decreasing or increasing the free energy of binding beyond the indicated minimum or maximum thresholds (x-axis; nominal p-value<0.05) respectively, stratified by position in the structure. Only mutations with confident free energy changes are shown (see Figure S19 for similar plots showing all mutations). **e.** Total numbers of mutations decreasing or increasing *bindingPCA* fitness (i.e. the fraction of bound protein complex) beyond the indicated minimum or maximum thresholds (x-axis) respectively. **f.** Similar to (**e**) except only mutations without significant effects on *abundancePCA* fitness are shown. See Figure S16 for the GB1 domain.

In addition to the average absolute change in the free energy of binding being higher within residues comprising the ‘sector’ defined by 20 coevolving residues in PSD95-PDZ3^10,51^ (two-sided Mann-Whitney U test p-value=1.5e-04, AUC=0.78), allosteric mutations themselves are highly enriched at these sites (odds ratio=8.3, two-sided Fisher’s Exact Test p-value<2.2e-16, Figure S17a), indicating that this network of physically proximal sites partially identifies the patterns of allostery described here. We also find that the probability that a mutation outside the binding interface will be allosteric significantly depends on its distance to the ligand in GRB2-SH3 and PSD-PDZ3 (Spearman’s ρ=−0.3 and −0.58 respectively, Figure 6c, Figure S16c), further suggesting the propagation of local perturbations to neighbouring residues as an important cause of allosteric effects observed in these domains. Also consistent with this, allosteric coupling scores estimated by a network-based perturbation propagation algorithm using only structural contacts^52^ correlate with the proportion of allosteric mutations per residue in both GRB2-SH3 and PSD95-PDZ3 (Spearman’s ρ=0.51 and 0.42 respectively, Figure S18).

If the disruption of local energetic couplings is indeed an important initiator of allosteric effects, we reasoned that the identities of the original and substituted residues in allosteric mutations should be enriched in specific AA types. Indeed, allosteric mutations are significantly enriched at Glycine residues in all three protein domains considered (odds ratio=2.4-5.7, two-sided Fisher’s Exact Test p-value<9e-5, Figure 6c and Figure S17b), whose replacement would increase the local mass, volume and also conformational rigidity. Glycine residues also comprise two major allosteric sites in each domain. In fact, across all three protein domains, four out of five Glycines that occur in secondary structure elements (and outside the binding interface) are major allosteric sites (odds ratio=20, two-sided Fisher’s Exact Test p-value=0.01, Figure 6c and Figure S17c). Likewise, changes to Proline are consistently the most enriched mutant AA in allosteric mutations (odds ratio=2.7-5.8, two-sided Fisher’s Exact Test p-value<2e-2, Figure 6c and Figure S17b) with this residue’s exceptional rigidity likely to introduce both local structural distortion and altered dynamics, both of which may be important for allosteric communication^7^.

### Mutational landscapes for network re-wiring

Disease-causing and evolutionarily-selected mutations have previously been conceptualised as perturbing cellular processes by altering either the ‘nodes’ or ‘edges’ of PPI networks^53,54^. However, systematic data quantifying the effects of mutations on network edges is limited to a small number of mutations for any individual protein^53,55^. We therefore used our data to further investigate the properties of ‘edgetic’^53,55^ network-altering mutations.

We first considered changes in binding affinity as the trait of interest. Mutations in ligand-contacting sites are very biased towards disruption (Figure 6d, Figure 3a-c, Figure S19) with many fewer mutations increasing rather than decreasing binding affinity. This is consistent with the binding interface residues being near optimal for this function. Mutations at ligand-distal surface positions tend to have milder effects on the free energy of binding than those at ligand-proximal sites, but their total number is greater and their direction less biased (Figure 6d, Figure S19). Indeed, the number of mutations at surface residues increasing binding affinity is greater than the number of disrupting mutations in the binding interface (110 vs. 91 and 143 vs. 126 for GRB2-SH3 and PSD95-PDZ3 respectively).

We next considered changes in the fraction of bound protein complex as the phenotype of interest i.e. regardless of whether the underlying biophysical mechanism involves a change in binding or folding energy or both (Figure 6e, Figure S16d). Mutations in the protein core frequently and strongly reduce the fraction of bound protein complex, with those in PSD95-PDZ3 far outnumbering mutations in the binding interface and surface of similar effect size. Finally, we considered mutations that alter binding affinity without a significant change in abundance (Figure 6f). These mutations are comparatively rare in the protein core and indeed appear to be totally absent in GRB2-SH3. Unsurprisingly, mutations in the binding interface have strongly disruptive effects on binding. However, mutations in solvent exposed surface positions are particular in that they are both numerous and, especially in PSD95-PDZ3, can fine tune binding affinity in both directions without disrupting stability.

In summary, the high density of allosteric mutations throughout these domains suggests a larger mutational target space for ‘edgetic’ network-altering genetic variants than has been previously appreciated: many mutations outside of interaction interfaces should be expected to alter not just protein stability but also the affinities of proteins for their interaction partners.

## Discussion

We have presented here a general approach—multidimensional mutagenesis—to infer the *in vivo* biophysical effects of mutations and used a specific implementation of it—*ddPCA*—to produce the first global maps of allosteric mutations for any proteins. The multidimensional mutagenesis *ddPCA* approach quantifies the effects of mutations on multiple phenotypes in multiple genetic backgrounds, which is an efficient experimental design to infer the underlying causal biophysical effects of mutations by fitting thermodynamic models. We fit these models using neural networks with parameters having biophysical meaning, allowing the use of fast and user-friendly open source platforms for machine learning such as TensorFlow^56^.

This approach of fitting additive thermodynamic models to mutation scanning data is conceptually similar to previous pioneering work inferring the energetic effects of mutations in regulatory elements from combinatorial mutants^57–59^ and antibody affinities using ligand titrations^60^. Combined with a diversity of selection methods^32,61^, multidimensional mutagenesis strategies including *ddPCA* should facilitate the rapid and comprehensive mapping of the *in vivo* biophysical effects of mutations and the generation of free energy landscapes for diverse macromolecules, interactions and pathways.

The comprehensive energetic and allosteric landscapes presented here provide a number of important insights into protein function and evolution. First, allosteric mutations are common and their frequency increases closer to binding interfaces, suggesting local propagation of perturbations as an important molecular mechanism. Second, allosteric mutations are strongly enriched for certain AA changes, with mutations at Glycine residues in secondary structure elements particularly likely to be allosteric and mutations to Proline also frequently having allosteric effects. Third, mutations are frequently pleiotropic, affecting both stability (PPI network node) and affinity (PPI network edge). Fourth, mutations in both protein cores and surfaces can tune stability and affinity, suggesting high evolvability for new regulatory mechanisms and diverse opportunities for the modulation of protein abundance and interactions via drug binding.

The application of *ddPCA* and related methods should help accelerate allosteric drug discovery by producing global allosteric maps for diverse target proteins. In particular, PPIs represent a very large but difficult to inhibit therapeutic target space and the systematic identification of allosteric sites may open up opportunities for inhibiting targets currently considered ‘undruggable’^14^. Moreover, systematic maps of spatial information transfer in proteins—and selection experiments that identify modifiers of this transfer^16,21,22^—should provide fundamental insights into the mechanisms that underlie allostery, elucidate how it evolves, and quantify the extent to which allostery varies within and between protein families.

Understanding, predicting and engineering the encoding of biophysical properties by amino acid sequences is one of the most fundamental problems in molecular biology. That such a central problem remains unresolved after decades of research is, we would argue, primarily due to a lack of systematic and unbiased data quantifying how changes in sequence alter the biophysical properties of proteins. For fundamental problems where very large quantitative datasets already exist, dramatic recent progress has been made using deep learning, allowing, for example, the accurate prediction of protein structures from sequence^62^. However, for many other core problems of molecular biology, suitably diverse and quantitative training datasets do not yet exist: we still need to generate them. A key advantage of a general method such as *ddPCA* is that it can be potentially used to quantify the effects of millions of mutations on the biophysical properties of thousands of proteins, allowing three of the fundamental encoding problems of biology – protein folding (sequence-to-stability), binding (sequence-to-affinity) and allostery to be addressed using massive-scale perturbation experiments. We envisage that such large, quantitative datasets will allow machine learning approaches to be effectively brought to bear on the generative functions of molecular biology, including predicting macromolecular stability, affinity, specificity and allostery from sequence. If successful, this combination of brute force experimentation and machine learning will usher in a new era of *predictive* molecular biology, where the biophysical properties of proteins can be accurately determined and engineered. Such predictive ability would open up unprecedented possibilities in industrial, agricultural and environmental biotechnology, and would revolutionise clinical genetics and the development of therapeutics.

## Materials and methods

### Media and buffers used

- LB: 10 g/L Bacto-tryptone, 5 g/L Yeast extract, 10 g/L NaCl. Autoclaved 20 min at 120°C.
- YPDA: 20 g/L glucose, 20 g/L Peptone, 10 g/L Yeast extract, 40 mg/L adenine sulphate. Autoclaved 20 min at 120°C.
- SORB: 1 M sorbitol, 100 mM LiOAc, 10 mM Tris pH 8.0, 1 mM EDTA. Filter sterilized (0.2 mm Nylon membrane, ThermoScientific).
- Plate mixture: 40% PEG3350, 100 mM LiOAc, 10 mM Tris-HCl pH 8.0, 1 mM EDTA pH 8.0. Filter sterilized.
- Recovery medium: YPD (20 g/L glucose, 20 g/L Peptone, 10 g/L Yeast extract) + 0.5 M sorbitol. Filter sterilized.
- SC –URA: 6.7 g/L Yeast Nitrogen base without amino acid, 20 g/L glucose, 0.77 g/L complete supplement mixture drop-out without uracil. Filter sterilized.
- SC –URA/MET/ADE: 6.7 g/L Yeast Nitrogen base without amino acid, 20 g/L glucose, 0.74 g/L complete supplement mixture drop-out without uracil, adenine and methionine. Filter sterilized.
- Competition medium: SC –URA/MET/ADE + 200 ug/mL methotrexate (BioShop Canada Inc., Canada), 2% DMSO.
- DNA extraction buffer: 2% Triton-X, 1% SDS, 100mM NaCl, 10mM Tris-HCl pH8, 1mM EDTA pH8.

### *ddPCA* plasmids construction

Three generic plasmids were constructed to be able to assay any protein of interest by *bindingPCA* or *abundancePCA*: the *bindingPCA* plasmid (pGJJ001), the *abundancePCA* plasmid (pGJJ045) and the mutagenesis plasmid (pGJJ055).

The first two plasmids were derived from plasmid pGD110 which carries two halves of the murine methotrexate-resistant DHFR (DHFR1,2 and DHFR3) with C-terminus (GGGGS)_4_ linker fusions under the expression of CYC promoters and a shared CYC terminator (URA3 cassette plasmid for yeast auxotrophic selection during the selection assays) as described^33^ plus a multiple cloning site. pGJJ001 had the same structure as pGD110 but with a barcode cloning site upstream of the CYC promoter driving expression of DHFR3 in case a barcode-variant association sequencing strategy was necessary. To construct the plasmid, pGD110 was amplified in 3 different fragments using primer pairs oGJJ001-oGJJ002, oGJJ003-oGJJ083 and oGJJ82-oGJJ016 (Table S1), which were then assembled by Gibson reaction (prepared in house) at 50°C for one hour. The *abundancePCA* plasmid pGJJ045 was constructed by Gibson assembly by substituting the CYC promoter driving expression of the half DHFR1,2 for the GPD promoter using primer pairs oGJJ47-oGD087 and oGJJ46-oGD089 (Table S1).

The generic mutagenesis plasmid (pGJJ055) was created to harbour a landing site with HindIII and AvrII restriction sites so that the CYC promoter and DHFR3 fused to any protein of interest could be cloned into it to perform error-prone PCR, nicking mutagenesis^63^ or other alternative mutagenesis strategies. It was derived from pUC19, to avoid future plasmid selection in yeast if not properly purified (not containing any yeast auxotrophic cassette). The final plasmid pGJJ055 was built in two cloning steps. Initially pUC19 was reduced in size (the lacZα fragment was deleted) to increase the efficiency of *E. coli* transformation and three synonymous mutations were introduced in the ampicillin resistance cassette (*bla* gene) to remove specific restriction sites. This resulting intermediate plasmid (pGJJ003) was obtained using three-fragment Gibson assembly. Two fragments were amplified (primer pairs oGJJ008-9 and oGJJ010-11) from pUC19 that introduced the barcode landing site. The third fragment with the *bla* sequence with the synonymous mutations was synthesized as a dsDNA gene block (gbGJJ001, GeneScript, Table S2). To generate the final mutagenesis plasmid (pGJJ055) a T>C synonymous substitution was added in the *bla* gene of pGJJ003 to create an Nb.BbvCI restriction site by one-fragment Gibson assembly using primers oGJJ049-50.

### GRB2-SH3 domain plasmid construction

To construct the GRB2-SH3 *abundancePCA* plasmid (pGJJ046) the 56 amino acid long SH3 domain of GRB2 (from amino acid 159 to 224 of the human protein GRB2) was fused to the C-terminus of the DHFR3 fragment of the *abundancePCA* plasmid (pGJJ045). To do so the SH3 domain was amplified by PCR reaction using primer pair oGJJ012-13 to introduce the flanking HindIII and NheI restriction sites and then cloned into the digested pGJJ045 plasmid using T4 Ligase (NEB).

To construct the GRB2-SH3 *bindingPCA* plasmid (pGJJ034), first the sequence of GAB2 containing the linear peptide (32 amino acids long, amino acid 498 to 530 of the human GAB2 protein) was fused to the fragment DHFR1,2. GAB2 was amplified using primer pair oGJJ014-15, which introduced flanking BamHI and SpeI restriction sites. Both the PCR product and the *bindingPCA* plasmid (pGJJ001) were digested and purified. The assembly of the new *bindingPCA* plasmid with GAB2 (pGJJ006) was obtained by ligation using T4 Ligase. After validation by Sanger sequencing, pGJJ006 was digested with HindIII and NheI restriction enzymes and cloned with the GRB2-SH3 domain to obtain the final wild-type GRB2-SH3 *bindingPCA* plasmid pGJJ034 (with both GRB2-SH3 and GAB2 linear peptide fused to both fragments of DHFR).

### PSD95 PDZ domain plasmid construction

To construct the PSD95-PDZ3 *abundancePCA* plasmid (pGJJ068) the sequence of the third PDZ domain of PSD95 (from amino acid 354 to 437 of the human DLG4 gene) was fused to the C-terminus of the DHFR3 fragment of the *abundancePCA* plasmid (pGJJ045). A gene block containing the PSD95-PDZ3 domain sequence was ordered (gbGJJ010, IDT, Table S2), amplified by PCR (Q5-Hot Start Polymerase, NEB) using oligos oGJJ078-79 to add a HindIII and NheI restriction sites, digested with HindIII and NheI and subsequently cloned into the digested pGJJ045 by T4 Ligase (NEB) standard protocol.

To obtain the *bindingPCA* plasmid of PSD95-PDZ3 (pGJJ072), the entire human gene CRIPT was fused to the DHFR1,2 half of the *bindingPCA* plasmid pGJJ001. To do so, a gene block with the sequence of the human CRIPT gene was ordered with flanking BamHI and SpeI restriction sites (gbGJJ011, IDT). The gene block was digested and cloned into pGJJ001, creating the intermediate plasmid pGJJ066. pGJJ066 was later digested with HindIII and NheI to clone in the C-terminus of the DHFR3 the digested PCR product of the PSD95-PDZ3 domain, obtaining the final wild-type PSD95-PDZ3 *bindingPCA* plasmid pGJJ072.

### Growth rate measurements of individual constructs

Seven single amino acid mutations of the GRB2-SH3 domain were tested for yeast growth in the presence of MTX on both abundance and binding assays. To construct these plasmids, 7 gene blocks containing the mutated versions of GRB2-SH3 were synthesized with HindIII and NheI flanking restriction sites (gbGJJ002-8, Twist Bioscience, Table S2). The gene blocks were digested, purified and assembled into the previously digested and purified pGJJ006 and pGJJ045 (*bindingPCA* plasmid with GAB2 and *abundancePCA* plasmid, respectively).

The GRB2-SH3 wild-type and mutant constructs were individually transformed into yeast following a small-scale high efficiency yeast transformation (See section 7.1, same protocol but scaled down in volume 0.0003X). After transformation, different colonies for each construct were picked and grown independently in 400 uL of SC –URA/MET/ADE overnight at 30°C using a 96 deep well plate. The following morning the optical density (OD) of each well was measured using a Tecan Infinite M Plex plate reader (Tecan, Switzerland). Cultures in each well were diluted to an OD_600nm_ of 0.1. For the selection experiment, 5 uL of the diluted cells were added into 95 ul of 1.053X competition media (SC –URA/MET/ADE + 200 ug/mL methotrexate) to obtain 100 uL of starting culture at OD_600nm_ = 0.005 of uL of per well. The culture was grown for 60h at 30°C in a Tecan plate reader where OD_600nm_ measurements were taken every 15 minutes. The growth rate in each well was obtained by calculating the slope of the exponential phase of the growth curve (slope of a linear fit of the log10(OD_600nm_) against time).

### Mutagenesis library construction

1. **Types of mutant libraries:** In this study two types of mutagenesis library construction approaches were used: error-prone PCR or one-pot nicking mutagenesis^63^. The selection of the technique depended on the domain length, the number of targeted mutations and the mutational biases of the strategies (mutation type vs. positional bias for error-prone PCR and nicking mutagenesis respectively). Table S3 contains fully detailed information of the different libraries constructed in this study. For the GRB2-SH3 domain two types of libraries were constructed (Table S3). An initial error-prone PCR library, covering some singles and few double mutants, which was used in the *bindingPCA* assay (“SH3_EP”), and a second library of two rounds of nicking mutagenesis, extending the amount of single and double mutants to run the *abundancePCA* assay (“SH3_NM2”). Given the short length of the GRB2-SH3 domain, these two high complexity libraries could be sequenced in a HiSeq 125bp paired-end sequencing platform. Due to the long length of the PSD95-PDZ3 domain, the platform for sequencing used was the MiSeq 250bp paired-end, giving a substantially lower number of reads counts per run compared to the HiSeq. Thus the PSD95-PDZ3 mutant libraries were constructed to have lower complexity compared to the SH3 ones. This consisted of a shallow double mutant library of one round of nicking mutagenesis, starting from a template pool of 10 different PSD95-PDZ3 wild-type and single mutant backgrounds (“PDZ_NM2sha”).
2. **Cloning the SH3 and PDZ domains into the mutagenesis plasmid** The CYC promoter and DHFR3 C-terminally fused to the GRB2-SH3 domain was cloned into the mutagenesis plasmid (pGJJ055) by digestion-ligation protocol. The CYC-DHFR3-GRB2 insert was obtained by digesting the plasmid pGJJ025 (*bindingPCA* plasmid with GRB2-SH3 tagged to DHFR3) with HindIII-HF and AvrII and purifying the correct size band using the QIAquick Gel Extraction Kit (QIAGEN). The mutagenesis plasmid pGJJ055 was digested with HindIII-HF and AvrII and purified with the MinElute PCR Purification Kit (QIAGEN). The GRB2 mutagenesis plasmid (pGJJ057) was assembled by a ligation reaction (T4 Ligase, New England Biolabs) following the manufacturer’s protocol. After transformation into NEB10-beta High Efficiency competent cells, the plasmid sequence was verified by Sanger Sequencing (GATC, Eurofins Genomics). The mutagenesis plasmid containing the PSD95-PDZ domain (pGJJ111) was obtained following the same strategy as for the GRB2-SH3 mutagenesis plasmid. The insert containing the CYC promoter and DHFR3 fused to the PDZ domain was digested and purified from plasmid pGJJ072 and subsequently cloned as described above into the HindIII-AvrII digested pGJJ055.
3. **SH3 error-prone PCR library** The error-prone PCR reaction for libraries “SH3_EP” was done using the GeneMorph II Mutagenesis Kit (Agilent Technologies) following the manufacturer’s protocol. Primers oGJJ048-152 (with homology to the origin of replication and the CYC promoter respectively) were used to amplify 1,011 bp of the SH3 (pGJJ057) mutagenesis plasmids. A single PCR reaction of 50 uL was run using 0.91 ng of template plasmid pGJJ057 (reaching the lowest recommended amount of plasmid by the manufacturer, lower plasmid amount increases the mutation frequency), with an annealing temperature of 56.4°C (previously determined by gradient PCR), 1 minute and 10 seconds of extension time. The PCR product was later run on an agarose gel for band confirmation and purified using the MinElute PCR Purification Kit (QIAGEN). The entire error-prone PCR product was digested with HindIII-HF and NheI-HF restriction enzymes. The correct band with the GRB2-SH3 domain was purified using the MinElute Gel Extraction Kit (QIAGEN) and the sample was quantified using a Qubit fluorometer to be cloned into the yeast assay plasmids by high efficiency temperature-cycle ligation^64^.
4. **GRB2-SH3 and PSD95-PDZ3 nicking mutagenesis libraries** The plasmid-based one-pot saturation (nicking) mutagenesis protocol was extracted from ^63^. The protocol was followed exactly as described, with two minor changes: the template plasmid DNA was prepared from 4°C midi-preps and the mutagenic primers were used at 1:10 molar ratio to plasmids instead of 1:20. The mutagenic primers were designed to contain the three degenerate bases (NNK) in the codon position with an average of 21bp of homology sequence upstream and downstream with the length adjusted to match similar melting temperatures (Sigma-Aldrich, ‘seq_opt’ column in Table S5). The mutant library “SH3_NM2” (Table S3) consisted of two rounds of nicking mutagenesis, using an equimolar mix of degenerate GRB2-SH3 primers (Table S5). In the first round of nicking mutagenesis, 3.3×10^5^ *E. coli* tranfromants were obtained. The resulting midiprep of the overnight culture was used as template for the second round of nicking mutagenesis with the same primer mix, but scaled up in volume 10 times. The second round of nicking mutagenesis resulted in 3.75×10^7^ *E. coli* transformants. For the “PDZ_NM2sha”, an initial single round of nicking mutagenesis using an equimolar mix of degenerated PSD95-PDZ3 primers (Table S5) was obtained for two reasons: (1) To obtain 9 random single mutants to use as template for another round of nicking mutagenesis (by randomly selecting single colonies and verified by Sanger sequencing) and (2) to quantify the degenerate primer positional bias and compensate for it in the shallow double mutant library by spiking part of this library was MiSeq 250bp (data not shown). To construct the final “PDZ_NM2sha” library an equimolar pool of nine single mutants and the wild-type was used as the plasmid template for a round of nicking mutagenesis. To compensate for the extreme positional biases, each mutagenic primer was mixed in the pool inversely to the mean read counts per position from the sequencing results of MiSeq spike-in initial library. The libraries midi-preps were digested with HindIII and NheI restriction enzymes and the insert containing the mutated protein was gel purified (MinElute Gel Extraction Kit, QIAGEN) to be later cloned into the two assay plasmids by temperature-cycle ligation.
5. **Cloning the mutant libraries into the yeast assay plasmids** Both, the *abundancePCA* plasmid (pGJJ045) and the *bindingPCA* plasmid containing the ligand GAB2 or CRIPT fused to DHFR1,2 (pGJJ006 and pGJJ066 respectively) were digested with the same enzymes and purified using the QIAquick Gel Extraction Kit (QIAGEN). The assembly of GRB2 SH3 and PSD95 PDZ mutant libraries in both assay plasmids were done overnight by temperature-cycle ligation using T4 ligase (New England Biolabs) according to the manufacturer's protocol, 67 fmol of backbone and 200 fmol of insert in a 33.3 uL reaction. The ligation was desalted by dialysis using membrane filters for 1h and later concentrated 3.3X using a SpeedVac concentrator (Thermo Scientific). All concentrated assembled libraries were transformed into NEB 10β High-efficiency Electrocompetent *E. coli* cells according to the manufacturer’s protocol (volumes used in each library specified in Table S3). Cells were allowed to recover in SOC medium (NEB 10β Stable Outgrowth Medium) for 30 minutes and later transferred to 200 mL of LB medium with ampicillin 4X overnight. The total number of estimated transformants for each library can be found in Table S3. 100 mL of each saturated *E. coli* culture were harvested next morning to extract the plasmid library using the QIAfilter Plasmid Midi Kit (QIAGEN).

### Methotrexate selection assays

1. **Large-scale yeast transformations** The high-efficiency yeast transformation protocol of all the different selection assays was derived from^33^. The protocol was scaled in volume depending on the targeted number of transformants of each library. The transformation protocol described below (adjusted to a pre-culture of 700 mL of YPDA) was scaled up or down in volume as reported in Table S3. For each of the two assays (*bindingPCA* and *abundancePCA*) of each protein domain library three independent pre-cultures of BY4742 were grown in 80 mL standard YPDA at 30°C overnight. The next morning, the cultures were diluted into 700 mL of pre-wormed YPDA at an OD_600nm_ = 0.3. The cultures were incubated at 30°C for 4 hours. After growth, the cells were harvested and centrifuged for 5 minutes at 3,000g, washed with sterile water and later with SORB medium (100mM LiOAc, 10mM Tris pH 8.0, 1mM EDTA, 1M sorbitol). The cells were resuspended in 34.4 mL of SORB and incubated at room temperature for 30 minutes. After incubation, 700 μL of 10mg/mL boiled salmon sperm DNA (Agilent Genomics) was added to each tube of cells, as well as 14 μg of plasmid library. After gentle mixing, cells were split in tubes of ~8.8 mL of cells and 35 mL of Plate Mixture (100mM LiOAc, 10mM Tris-HCl pH 8, 1mM EDTA/NaOH, pH 8, 40% PEG3350) were added to each tube to be incubated at room temperature for 30 more minutes. 3.5 mL of DMSO was added to each tube and the cells were then heat shocked at 42°C for 20 minutes (inverting tubes from time to time to ensure homogenous heat transfer). After heat shock, cells were centrifuged and re-suspended in ~50 mL of recovery media and allowed to recover for 1 hour at 30°C. Next, cells were again centrifuged, washed with SC-URA medium and re-suspended in SC –URA (volume used in each library found in Table S3). After homogenization by stirring, 10 uL were plated on SC –URA Petri dishes and incubated for ~48 hours at 30°C to measure the transformation efficiency. The independent liquid cultures were grown at 30°C for ~48 hours until saturation. The number of yeast transformants obtained in each library assay can be found in Table S3.
2. **Selection assays** For each of the *bindingPCA* or *abundancePCA* assays, each of the growth competitions was performed right after yeast transformation. After the first cycle of post-transformation plasmid selection, a second plasmid selection cycle (input) was performed by inoculating SC –URA/MET/ADE at a starting OD_600nm_ = 0.1 with the saturated culture (volume of each experiment specified in Table S3). Cells were grown for 4 generations at 30°C under constant agitation at 200 rpm. This allowed the pool of mutants to be amplified and enter the exponential growth phase. The competition cycle (output) was then started by inoculating cells from the input cycle into the competition media (SC –URA/MET/ADE + 200 ug/mL Methotrexate) so that the starting OD_600nm_ was 0.05. For that, the adequate volume of cells were collected, centrifuged at 3,000 rpm for 5 minutes and resuspended in the pre-warmed output media. Meanwhile, each input replicate culture was splitted in two and harvested by centrifugation for 5 min at 5,000g at 4°C. Yeast cells were washed with water, pelleted and stored at −20°C for later DNA extraction. After ~5-5.2 generations of competition cycle, each output replicate culture was splitted into two and harvested by centrifugation for 5 min at 5,000g at 4°C, washed twice with water and pelleted to be stored at −20°C.

### DNA extractions and plasmid quantification

The DNA extraction protocol used is described below for a 400 mL harvested culture of OD_600nm_ ~ 1.6. Depending on the volume harvested in each selection assay (Table S3), the following protocol was scaled up or down in volume.

Cell pellets (one for each experiment input/output replicate) were re-suspended in 4 mL of DNA extraction buffer, frozen by dry ice-ethanol bath and incubated at 62°C water bath twice. Subsequently, 4 mL of Phenol/Chloro/Isoamyl 25:24:1 (equilibrated in 10mM Tris-HCl, 1mM EDTA, pH8) was added, together with 4 g of acid-washed glass beads (Sigma Aldrich) and the samples were vortexed for 10 minutes. Samples were centrifuged at RT for 30 minutes at 4,000 rpm and the aqueous phase was transferred into new tubes. The same step was repeated twice. 0.4 mL of NaOAc 3M and 8.8 mL of pre-chilled absolute ethanol were added to the aqueous phase. The samples were gently mixed and incubated at −20°C for 30 minutes. After that, they were centrifuged for 30 min at full speed at 4°C to precipitate the DNA. The ethanol was removed and the DNA pellet was allowed to dry overnight at RT. DNA pellets were resuspended in 2.4 mL TE 1X and treated with 20 uL of RNaseA (10mg/mL, Thermo Scientific) for 30 minutes at 37°C. To desalt and concentrate the DNA solutions, QIAEX II Gel Extraction Kit was used (200 μL of QIAEX II beads). The samples were washed twice with PE buffer and eluted 500 μL of 10 mM Tris-HCI buffer, pH 8.5. Finally, plasmid concentrations in the total DNA extract (that also contained yeast genomic DNA) were quantified by qPCR using the primer pair oGJJ152-oGJJ153, that bind to the ori region of the plasmids.

### Sequencing library preparation

The sequencing libraries were constructed in two consecutive PCR reactions. The first PCR (PCR1) was designed to amplify the mutated protein of interest and to increase the nucleotide complexity of the first sequenced bases by introducing frame-shift bases between the adapters and the sequencing region of interest. The second PCR (PCR2) was necessary to add the remainder of the Illumina adapter and demultiplexing indexes.

To avoid PCR biases, PCR1 of each independent sample (input/output replicates of any of the yeast assays) was run with an excess of plasmid template 20-50 times higher than the number of expected sequencing reads per sample (Table S3). Each reaction started with a maximum of 1.25×10^7^ template plasmid molecules per uL of PCR1, avoiding introducing more yeast genomic DNA that interfered with the efficiency of the PCR reaction. For this reason, PCR1s were scaled up in volume as specified in Table S3. The PCR1 reactions were run using Q5 Hot Start High-Fidelity DNA Polymerase (New England Biolabs) according to the manufacturer’s protocol, with 25 pmol of pooled frame-shift primers oGJJ52/54-58 and oGJJ84-89 (forward and reverse primers were independently pooled according to the nucleotide diversity of each oligo, Table S1). The PCR reactions were set to 60°C annealing temperature, 10 seconds (for GRB2-SH3 libraries) or 15 seconds (for PSD95-PDZ3 libraries) of extension time and run for 15 cycles. Excess primers were removed by adding 0.04 uL of ExoSAP-IT (Affymetrix) per uL of PCR1 reaction and incubated for 20 min at 37 ̊C followed by an inactivation for 15 min at 80 ̊C. The PCRs of each sample were then pooled and purified using the MinElute PCR Purification Kit (QIAGEN) according to the manufacturer’s protocol. DNA was eluted in EB to a volume 6 times lower than the total volume of PCR1.

PCR2 reactions were run for each sample independently using Hot Start High-Fidelity DNA Polymerase. The total reaction of PCR2 was reduced to half of PCR1, using 0.05 uL of the previous purified PCR1 per uL of PCR2. In this second PCR the remaining parts of the Illumina adapters were added to the library amplicon. The forward primer (5’ P5 Illumina adapter) was the same for all samples, while the reverse primer (3’ P7 Illumina adapter) differed by the barcode index (oligo sequences in Table S1), to be subsequently pooled together and demultiplex them after deep sequencing (indexes used in each replicate of each sequencing run found in Table S4). 10 cycles of PCR2s were run at 62°C of annealing temperature and 15 seconds (for GRB2-SH3 libraries) or 20 seconds (for PSD95-PDZ3 libraries) of extension time. All reactions from the same sample were pooled together and an aliquot was run on a 2% agarose gel to be quantified. After quantification, samples with different Illumina indexes that were sequenced together in the same flow-cell were pooled in an equimolar ratio, run on a gel and purified using the QIAEX II Gel Extraction Kit. The purified amplicon library pools were subjected to Illumina 125bp paired-end HiSeq (SH3 libraries) or 250bp paired-end MiSeq (PDZ libraries) sequencing at the CRG Genomics Core Facility.

### Sequencing data processing

FastQ files from paired-end sequencing of all *bindingPCA* and *abundancePCA* experiments were processed with DiMSum v1.2.8^39^ using default settings with minor adjustments:https://github.com/lehner-lab/DiMSum. All experimental design files and bash scripts with command-line options required for running DiMSum on these datasets are available on GitHub (https://github.com/lehner-lab/doubledeepmsx). In all cases, adaptive minimum Input read count thresholds based on the corresponding number of nucleotide substitutions (“fitnessMinInputCountAny” option) were selected in order to minimise the fraction of reads per variant related to sequencing error-induced “variant flow” from lower order mutants^39^.

For the PSD95-PDZ3 data, where nicking mutagenesis was performed on a starting template pool of 10 different backgrounds, variant counts associated with all samples (output from DiMSum stage 4) were further filtered using a custom script to retain double AA variants consisting of single AA mutations in these designed backgrounds. All read counts associated with remaining double AA variants (likely the result of PCR and sequencing errors) were discarded. Finally, fitness estimates and associated errors were then obtained from the resulting filtered variant counts with DiMSum (“countPath” option).

The pre-processed DMS data for GB1 were obtained from the supplementary information of a previous study^28^ (github.com/jotwin/ProteinGthermo). The data consist of sequencing read counts from a single (common) Input sample and three replicate (selection) experiments assaying the binding affinity of wild-type, single and double AA GB1 variants to IgG (Output samples)^44^. Fitness estimates and associated errors were obtained from variant counts with DiMSum v1.2.8 using default settings.

### Protein fitness estimation

For DMS experiments in general, the enrichment score (ES) for each variant *i* in replicate *r* is typically defined as the ratio between its frequency before and after selection:

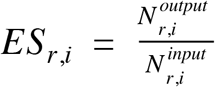

where *N*^*input*^ *N*^*output*^ are the corresponding numbers of sequenced reads in the Input and Output sample respectively. DiMSum calculates fitness scores for each variant *i* in replicate *r* as the natural logarithm of the enrichment score normalised to the wild-type or reference variant (*wt*):

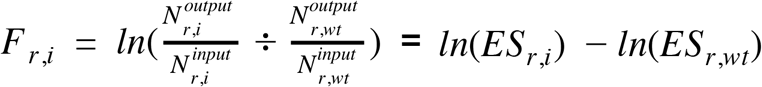

Measurement errors of fitness scores are estimated by fitting a modified Poissonian (count-based) model to the data—with replicate-specific multiplicative and additive terms that are common to all variants—that has been shown to outperform previous methods in benchmarking analyses^39^. Final DiMSum fitness and error estimates for all variants are obtained by merging between replicates using weighted averaging.

For cell growth based assays such as *bindingPCA* and *abundancePCA*, the growth rate associated with variant *i* in replicate *r* can be estimated as:

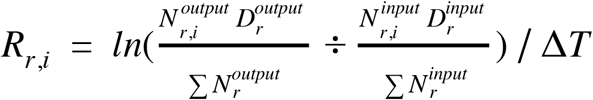

where *D*^*input*^ and *D*^*output*^ are proportional to the total number of cells in the Input and Output sample respectively (i.e. optical density or similar) and **Δ*T*** is the time difference (in hours) between measurements (i.e. selection/competition time). Growth rates and fitness scores can therefore be seen to be related by a simple linear transformation:

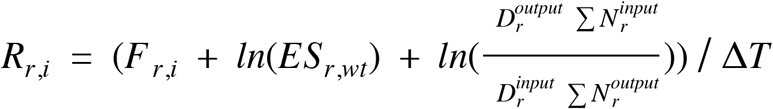

because all terms on the right-hand side are constant for all variants *i* in replicate *r*, except the fitness score *F*_*r,i*_.

### Thermodynamic models

At thermal equilibrium, the Boltzmann distribution relates the probability that a system will be in a given state *i* to the (Gibbs) free energy *G*_*i*_ of the state and the temperature of the system *T*:

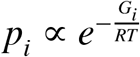

where *R* is the gas constant. Applied to proteins and considering the universe of *M* distinct conformations and/or interactions, the fraction of molecules in state *i* compared to all possible states is therefore given by:

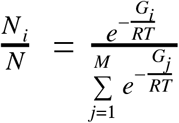

In the case of a two-state unfolded/folded model, where we denote the sum of energies of all possible unfolded states with the reference value of zero (i.e. *G*_*u*_ = 0), the fraction of molecules in the folded state is then:

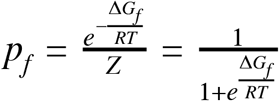

where 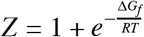 and Δ*G*_*f*_ is the energy difference between folded and unfolded states. As in previous work^28^, in order to capture changes in both fold stability and the stability of binding, we model PPIs as an equilibrium between three states: unfolded and unbound (*uu*), folded and unbound (*fu*), and folded and bound (*fb*). The probability of the unfolded and bound state (*ub*) is assumed to be negligible. The probability of the folded and bound state is then given by:

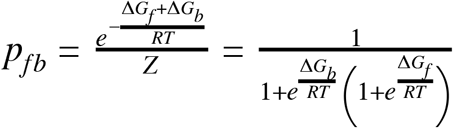

where 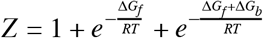 and Δ*G*_*b*_ is the energy difference between bound and unbound folded states (i.e. binding affinity). Importantly, both Δ*G*_*b*_ and Δ*G*_*f*_ implicitly depend upon the amino acid sequence. Furthermore, we assume that these energies are additive, meaning that the total free energy change (ΔΔ*G*) of an arbitrary variant *i* (of any mutational order) relative to the wild-type sequence is simply the sum over residue-specific energies (ΔΔ*g*) corresponding to all constituent individual (i.e. lowest order) amino acid changes:

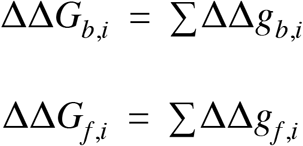

where ΔΔ*g*_*b,i*_ and ΔΔ*g*_*f.i*_ denote binding and folding free energy changes of constituent single amino acid substitutions of variant *i* relative to the wild-type. We can therefore express the absolute (rather than relative) free energies of binding and folding of an arbitrary variant *i* as:

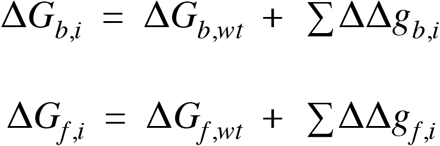

where Δ*G*_*b,wt*_ and Δ*G*_*f,wt*_ are the binding and folding free energies of the wild-type.

Considering that the concentration of the interaction partners represents a constant added to Δ*G*_*b*_, we expect this not to affect the estimation of *changes* in the free energy of binding (ΔΔ*G*_*b*_). Additionally, the effect (if any) of the fused DHFR fragments on the stability of the assayed protein domains Δ*G*_*f*_ is most likely additive in free energy space for all variants and similarly this would not affect the estimation of *changes* in the free energy of folding (ΔΔ*G*_*f*_). Finally, in the absence of calibrations to a reference set of directly measured free energies, our inferences should be strictly considered ‘pseudo’ free energy changes, but we refer to them without this qualifier in the interest of brevity.

### Thermodynamic model inference with MoCHI

Given that protein fitness (as defined above) is proportional to the concentration of either the free protein or protein complex in the *bindingPCA* or *stabilityPCA* assays respectively, we can parameterize binding fitness *F*_*fb*_ and abundance fitness *F*_*f*_ as follows:

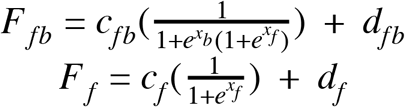

where *x*_*b*_ = Δ*G*_*b*_/*RT* and *x*_*f*_ = Δ*G*_*f*_/*RT* are additive functions of the amino acid sequence, and *c*_*fb*_, *c*_*f*_, *d*_*fb*_, *d*_*f*_ are scalar constants.

We trained a feed-forward neural network representation of this genotype-phenotype model using the backpropagation-based algorithms provided by TensorFlow 2^56^. We thereby inferred the binding and folding free energy changes that optimally predict *abundancePCA* and *bindingPCA* fitness measurements from the corresponding AA sequences. We refer to this method that infers the underlying biophysical traits from DMS data as MoCHI (in preparation).

The neural network architecture consists of one input layer (AA sequences) and one trainable hidden layer (additive traits i.e. free energies) for each biophysical trait (binding and folding), as well as one trainable hidden layer (linear transformation) per phenotype of interest (*abundancePCA* and *bindingPCA* fitness). An additional input layer specifies the corresponding phenotype associated with each sequence (Figure S2). Finally, the network includes two additional untrainable hidden layers to transform energies to bound and folded fractions respectively, as well as (helper) layers to multiplex/concatenate fitness predictions for different phenotypes into a single 1-dimensional output.

This architecture, i.e. a single 1-dimensional target (output) variable representing both *abundancePCA* and *bindingPCA* fitness, allows arbitrary combinations of variants for each phenotype to be used for training and also obviates the need to explicitly pair phenotype measurements of each variant.

The target (output) data to fit the neural network comprises fitness estimates for wildtype, single and double AA substitution variants. A random 10% of double AA substitution variants were held out and represent the validation data (unseen during training). Furthermore, in order to capture uncertainty in *ddPCA* fitness estimates, the training data was expanded 10x by randomly sampling ten values from the fitness error distribution of each variant. The validation data was left unchanged. In order to prioritise fitting of wild-type free energy terms (which influence final free energy estimates of all other variants) the wild-type variant was artificially duplicated to represent 1% of training data samples. This approach is conceptually similar to the practice of applying weights to important samples, which are used for weighting the loss function during training.

Input (feature) data comprises 1-hot encoded representations of the observed input sequences (wild-type, single and double AA substitutions) i.e. one binary feature variable for each unique AA substitution and each biophysical trait. An additional constant (bias term) was included for all variants to enable explicit fitting of the corresponding wild-type free energy terms (Δ*G*_*b,wt*_ and Δ*G*_*f,wt*_). Separate input layers for each biophysical trait enable their dimensions to be adjusted independently allowing distinct subsets of free energy terms to be inferred for binding and folding. Relatedly, we removed feature variables corresponding to free energy terms that are impossible to infer e.g. the free energy of binding for a given variant in the absence of this phenotype.

All models were trained for 1000 epochs using the Adam optimization algorithm with a learning rate of 0.001. The batch size hyperparameter was optimized by running a parameter sweep ([128, 256, 512, 1024]) and selecting the corresponding value with the lowest validation loss after 100 training epochs.

Free energies are calculated directly from model parameters as follows: Δ*G*_*b*_ = *x*_*b*_*RT* and Δ*G*_*f*_ = *x*_*f*_*RT*, where *T* = 303*K* and *R* = 0.001987. We estimated the confidence intervals of model-inferred free energies using a Monte Carlo simulation approach. The variability of inferred free energy changes was calculated between ten separate models fit using data from [1] independent random samples of fitness estimates from their underlying error distributions and [2] independent random samples of the validation data consisting of 10% of double AA substitution variants held out during training. Confident inferred free energy changes are defined as those with Monte Carlo simulation derived 95% confidence intervals <1kcal/mol (Figure S20).

## Data availability

All DNA sequencing data have been deposited in the Gene Expression Omnibus with accession number GSE184042.

## Code availability

Source code for fitting thermodynamic models (MoCHI), downstream analyses and to reproduce all figures described here is available at https://github.com/lehner-lab/doubledeepms.

## Acknowledgements

This work was funded by European Research Council (ERC) Advanced (883742) and Consolidator (616434) grants, the Spanish Ministry of Science and Innovation (BFU2017-89488-P, EMBL Partnership, Severo Ochoa Centre of Excellence), the Bettencourt Schueller Foundation, the AXA Research Fund, Agencia de Gestio d’Ajuts Universitaris i de Recerca (AGAUR, 2017 SGR 1322), and the CERCA Program/Generalitat de Catalunya. JMS was supported by an EMBO Long-Term Fellowship (ALTF 857-2016) and a Marie Skłodowska-Curie Fellowship (752809, EU Commission Horizon 2020).

## Author contributions

J.D., J.M.S., G.D. and B.L. conceived the project and designed the experiments; J.D., J.M.S. and C.H. performed the experiments; A.J.F. led the data analysis with help from J.D. and J.M.S.; A.J.F., J.M.S and B.L. formulated the thermodynamic model; A.J.F. wrote the code to implement and fit the model; B.L., A.J.F. and J.D. wrote the manuscript with input from all authors.

## Supplementary Figures

**Figure S1**. Number of doubles per single mutant and correlation between replicate read counts.

**Figure S2**. Neural network architecture.

**Figure S3**. Performance of thermodynamic models.

**Figure S4**. Performance of thermodynamic models after restricting the fitness data to a single phenotype or a single genetic background.

**Figure S5**. Performance of thermodynamic models after ddPCA data downsampling.

**Figure S6**. Comparisons of inferred free energy changes to smaller-scale datasets of previously reported in vitro measurements.

**Figure S7**. Binding and folding free energy landscapes of the GB1 domain.

**Figure S8**. Biophysical mechanism of mutations that affect binding.

**Figure S9**. GB1 mutational effects on protein stability.

**Figure S10**. Evolutionary conservation and hydrophobicity of surface de-stabilising residues.

**Figure S11**. Surface de-stabilising residues are involved in extra-domain interactions.

**Figure S12**. Major allosteric sites in the GB1 domain.

**Figure S13**. Changes in free energy of binding in ligand binding interfaces.

**Figure S14**. Changes in fitness and free energy of binding and folding of major allosteric sites.

**Figure S15**. Changes in fitness and free energy of binding and folding of allosteric sites and positions with allosteric mutations.

**Figure S16**. Allosteric mutations in GB1.

**Figure S17**. Enrichment of allosteric mutations in literature allosteric networks and specific residue types and classes.

**Figure S18**. Comparison of computationally predicted allosteric coupling scores to percentage of allosteric mutations per residue.

**Figure S19**. Mutational biases towards increased or decreased binding given the position in the domain structure.

**Figure S20**. Determining confident inferred free energy changes using a Monte Carlo simulation approach.

## Supplementary Tables

**Table S1.** Primers used in this study.

**Table S2.** Gene blocks used in this study.

**Table S3.** Experimental details and numbers of the mutagenesis libraries in this study.

**Table S4.** Illumina indexed primers combinations used in this study to demultiplex samples after deep sequencing.

**Table S5.** Degenerate NNK oligos used for the GRB2-SH3 and PSD95-PDZ3 nicking mutagenesis libraries.

**Table S6.** Fitness estimates for GB1, GRB2-SH3 and PSD95-PDZ3.

**Table S7.** Inferred folding and binding free energy changes and associated annotations for GB1, GRB2-SH3 and PSD95-PDZ3.

