## Supplementary figures for "Global mapping of the energetic and allosteric landscapes of protein binding domains"


Figure S1. Number of doubles per single mutant and correlation between replicate read counts. 3

Figure S2. Neural network architecture. 4

Figure S3. Performance of thermodynamic models. 6

Figure S4. Performance of thermodynamic models after restricting the fitness data to a single phenotype or a single genetic background. 7

Figure S5. Performance of thermodynamic models after ddPCA data downsampling. 9

Figure S6. Comparisons of inferred free energy changes to smaller-scale datasets of previously reported in vitro measurements. 10

Figure S7. Binding and folding free energy landscapes of the GB1 domain. 12

Figure S8. Biophysical mechanism of mutations that affect binding. 14

Figure S9. GB1 mutational effects on protein stability. 15

Figure S10. Evolutionary conservation and hydrophobicity of surface de-stabilising residues. 16

Figure S11. Surface de-stabilising residues are involved in extra-domain interactions. 17

Figure S12. Major allosteric sites in the GB1 domain. 19

Figure S13. Changes in free energy of binding in ligand binding interfaces. 21

Figure S14. Changes in fitness and free energy of binding and folding of major allosteric sites. 22

Figure S15. Changes in fitness and free energy of binding and folding of allosteric sites and positions with allosteric mutations. 23

Figure S16. Allosteric mutations in GB1. 25

Figure S17. Enrichment of allosteric mutations in literature allosteric networks and specific residue types and classes. 27

Figure S18. Comparison of computationally predicted allosteric coupling scores to percentage of allosteric mutations per residue. 28

Figure S19. Mutational biases towards increased or decreased binding given the position in the domain structure. 30

Figure S20. Determining confident inferred free energy changes using a Monte Carlo simulation approach. 32

Supplementary References 33

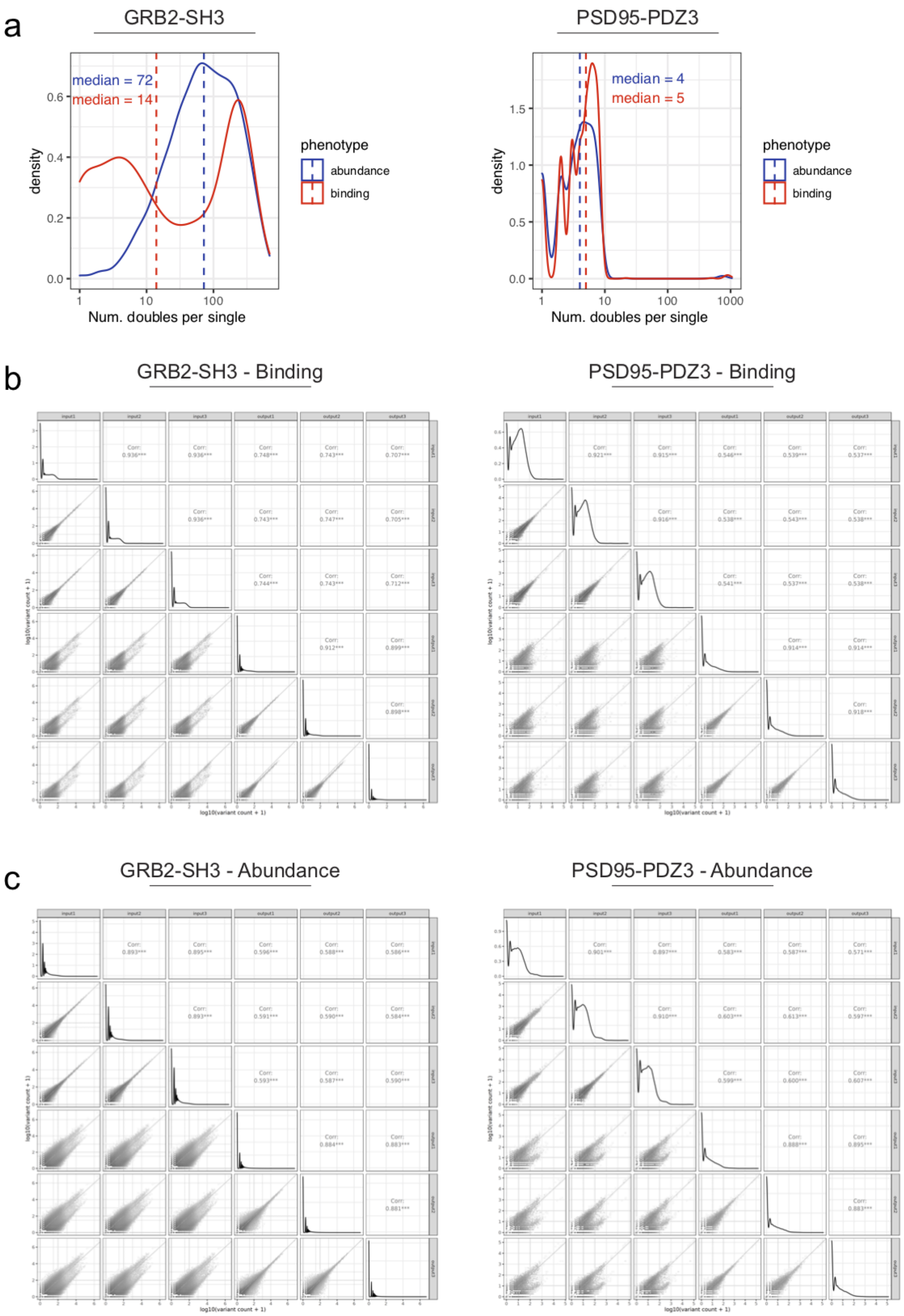
a
GRB2-SH3
median = 72
median = 14
density
Num. doubles per single
phenotype
abundance
binding
PSD95-PDZ3
median = 4
median = 5
density
Num. doubles per single
phenotype
abundance
binding
b
GRB2-SH3 - Binding
PSD95-PDZ3 - Binding
c
GRB2-SH3 - Abundance
PSD95-PDZ3 - Abundance

**Figure S1. Number of doubles per single mutant and correlation between replicate read counts.**

**a.** Distribution of the number of double AA substitutions comprising the same single AA substitution in the *abundancePCA* (blue) or *bindingPCA* (red) assays for the GRB2-SH3 (left) and PSD95-PDZ3 (right) protein domains. Median indicated with a dashed line and text label. **b-c.** Scatterplot matrices comparing read counts (and the addition of a pseudocount for log transformation) between all input and output samples of the *abundancePCA* (**b**) or *bindingPCA* (**c**) assay of the GRB2-SH3 (left) and PSD95-PDZ3 (right) domains. Corr=Pearson correlation.

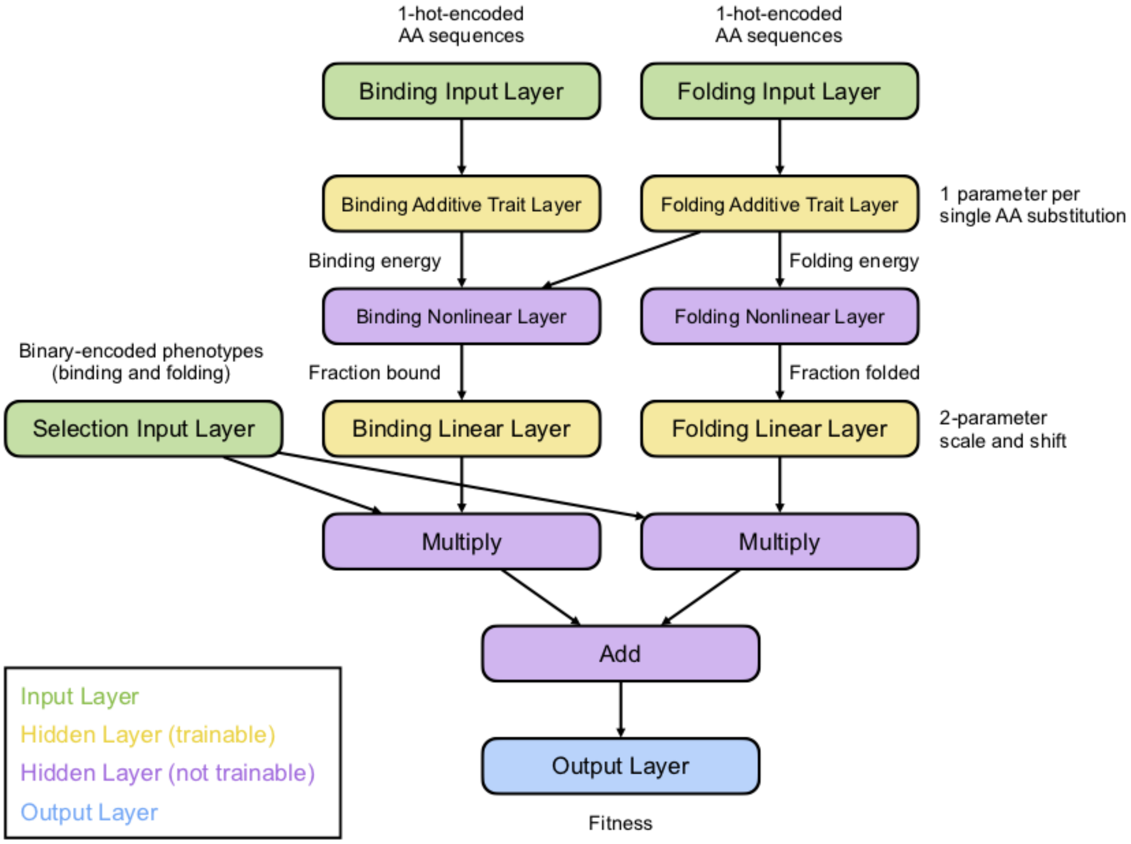


**Figure S2. Neural network architecture.**

Diagram of the neural network trained using TensorFlow in order to infer the binding and folding free energy changes that optimally predict *abundancePCA* and *bindingPCA* fitness measurements from 1-hot encoded representations of the corresponding input AA sequences.

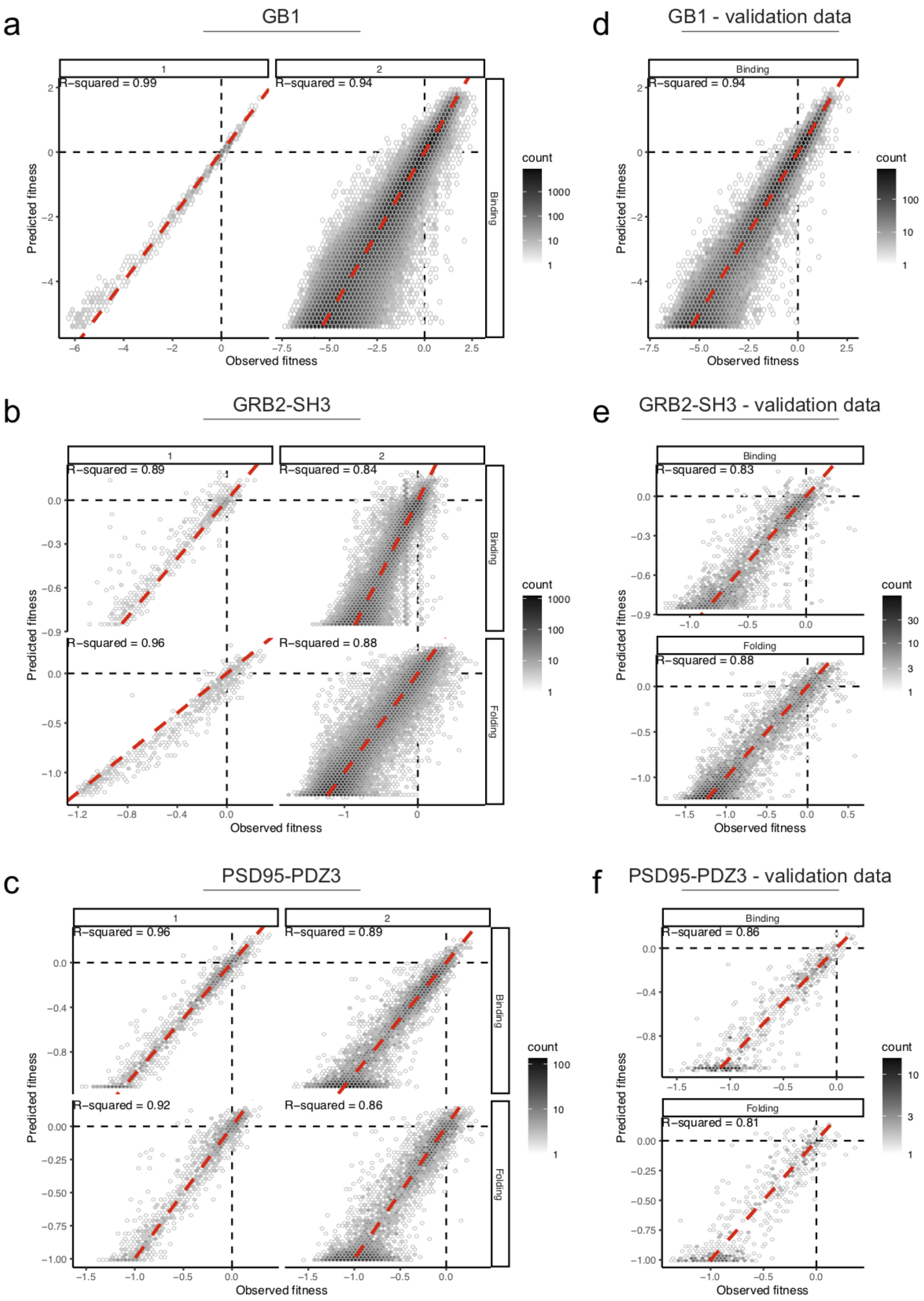

a
GB1
1
2
R−squared = 0.99
R−squared = 0.94
Binding
Predicted fitness
Observed fitness
count
d
GB1 - validation data
Binding
R−squared = 0.94
b
GRB2-SH3
R−squared = 0.89
R−squared = 0.84
R−squared = 0.96
R−squared = 0.88
Folding
e
GRB2-SH3 - validation data
R−squared = 0.83
R−squared = 0.88
c
PSD95-PDZ3
R−squared = 0.96
R−squared = 0.89
R−squared = 0.92
R−squared = 0.86
f
PSD95-PDZ3 - validation data
R−squared = 0.86
R−squared = 0.81

**Figure S3. Performance of thermodynamic models.**

**a-c.** 2d density plots comparing the *ddPCA* observed fitness and the model predicted fitness of single (left panels) and double AA substitutions (right panels) for the binding (top panels) and when existing, folding assays (bottom panels) of the GB1 (**a**), GRB2-SH3 (**b**) and PSD95-PDZ3 (**c**) domains. R-squared=Percentage of fitness variance explained. **d-f**. Same as (**a-c**) but using validation data comprising 10% of double mutants left out during model training.

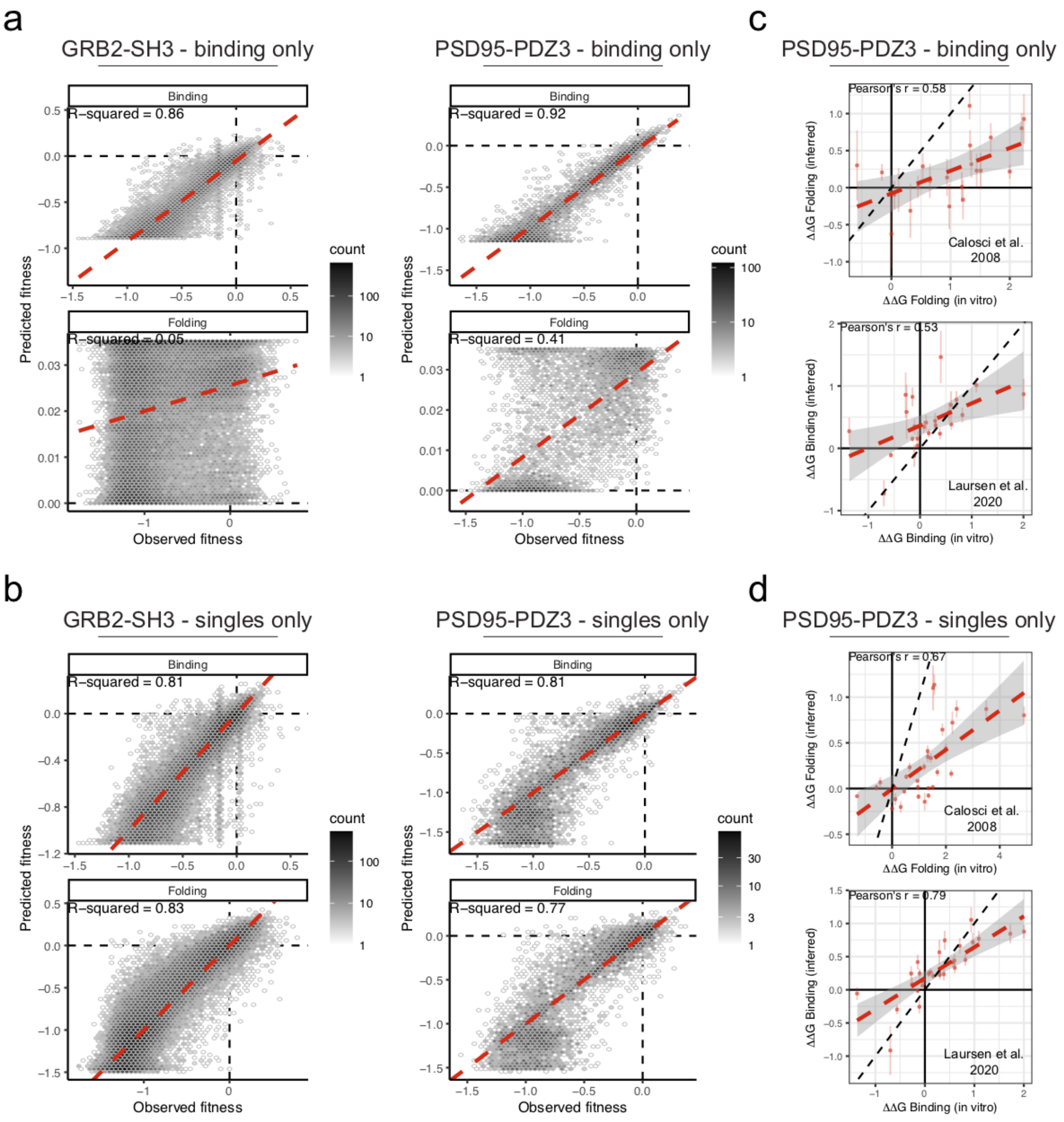


**Figure S4. Performance of thermodynamic models after restricting the fitness data to a single phenotype or a single genetic background.**

**a.** 2d density plots comparing the observed and predicted fitness of the binding (top panels) and abundance (bottom panels) assays when only the *bindingPCA* data is used for training the model for the GRB2-SH3 (left panels) and PSD95-PDZ3 (right panels). **b.** Same as in (**a**), but only using single mutant data from both binding and abundance assays to fit the models. R-squared=Percentage of fitness variance explained. **c-d.** Comparisons of inferred free energy changes to previously reported PSD95-PDZ3 mutant *in vitro* measurements where only *bindingPCA* data (**c**) or single mutants (**d**) were used to fit thermodynamic models. r=Pearson correlation.

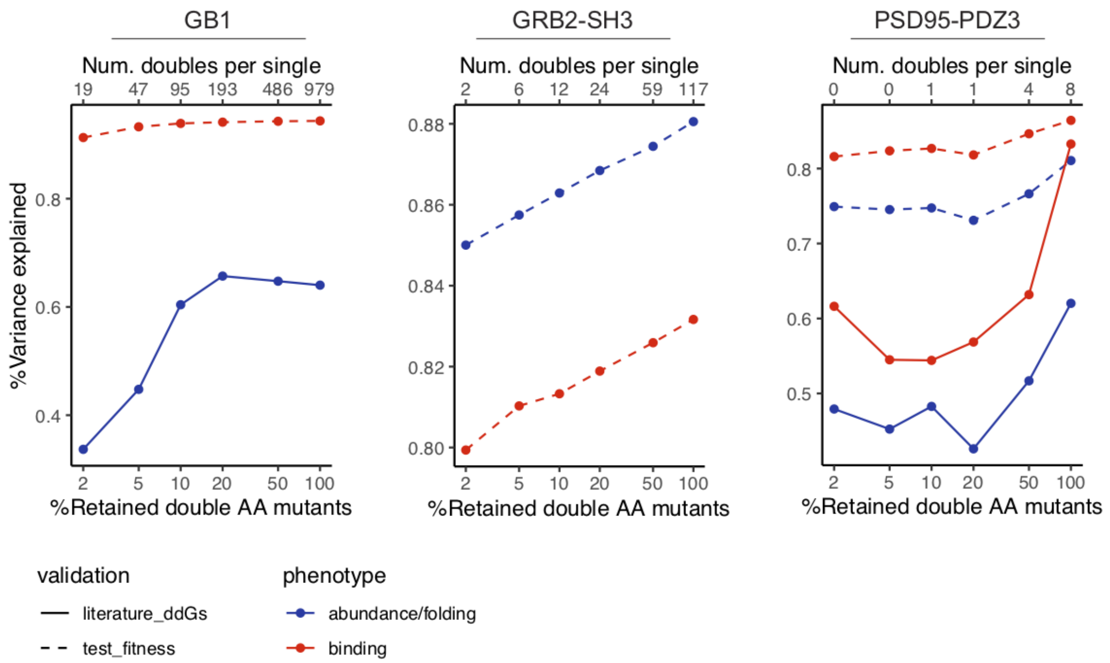


**Figure S5. Performance of thermodynamic models after *ddPCA* data downsampling.**

Dashed lines indicate the relationship between the percentage of fitness variance explained by model predictions with respect to held out validation data (10% of doubles) and the percentage of randomly retained double AA mutants used to train the model in the abundance (blue) or binding (red) assay. Results are shown separately for all protein domains. Solid lines indicate the relationship between the percentage variance explained by inferred free energies with respect to previously reported in vitro measurements for GB1 (Nisthal *et al.* 2019[1]) and PSD95-PDZ3 (Laursen *et al*. 2020[2] for ΔΔG binding, red; Calosci *et al*. 2008[3] for ΔΔG folding, blue), where models were trained using varying fractions of randomly downsampled double mutants (x-axis). The top scale indicates the median number of double AA mutants per single AA mutant in the full dataset.

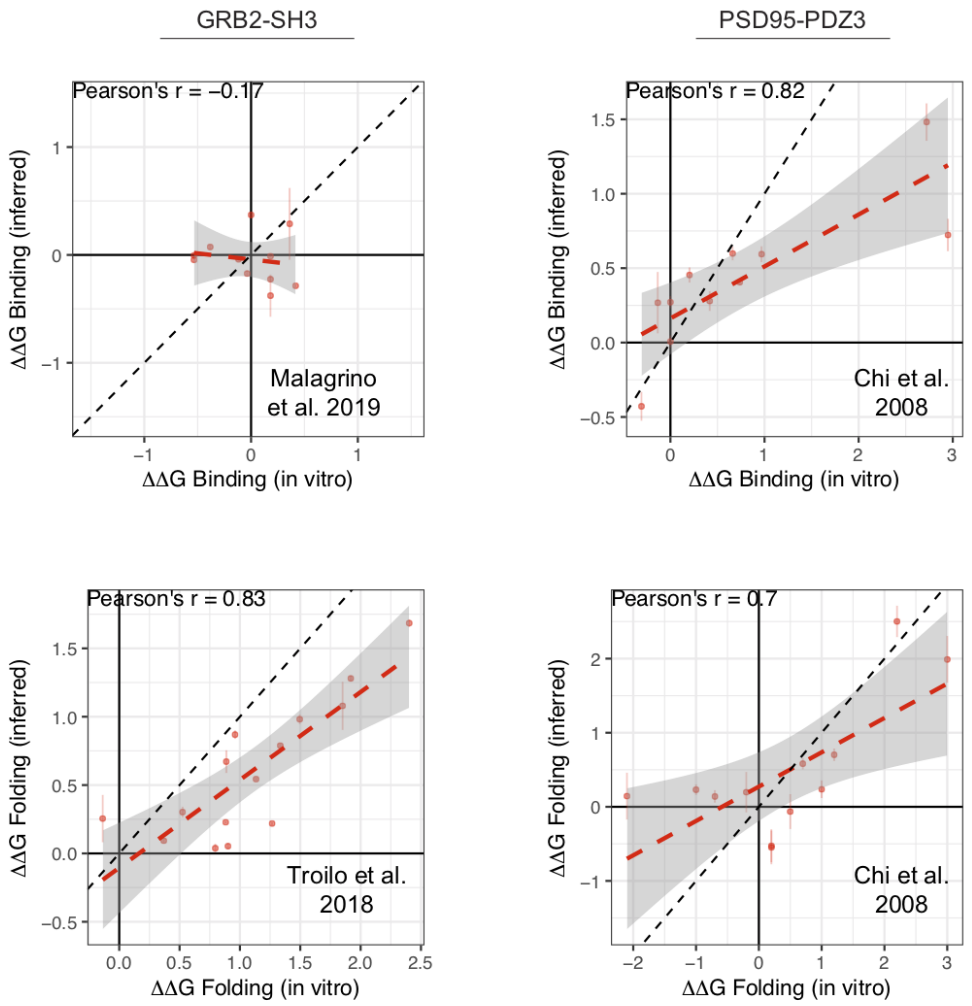


**Figure S6. Comparisons of inferred free energy changes to smaller-scale datasets of previously reported in vitro measurements.**

Comparisons of the model-inferred free energy changes to previously reported *in vitro* measurements for GRB2-SH3 (Malagrino *et al.* 2019[4] for ΔΔG binding and Troilo *et al.* 2018[5] for ΔΔG folding) and PSD95-PDZ3 (Chi *et al*. 2008[6]). Note the modest effect sizes of variants assayed in Malagrino *et al.* 2019. R=Pearson correlation. Inferred free energy changes' error bars indicate 95% confidence intervals (*in vitro* error measurement not provided).

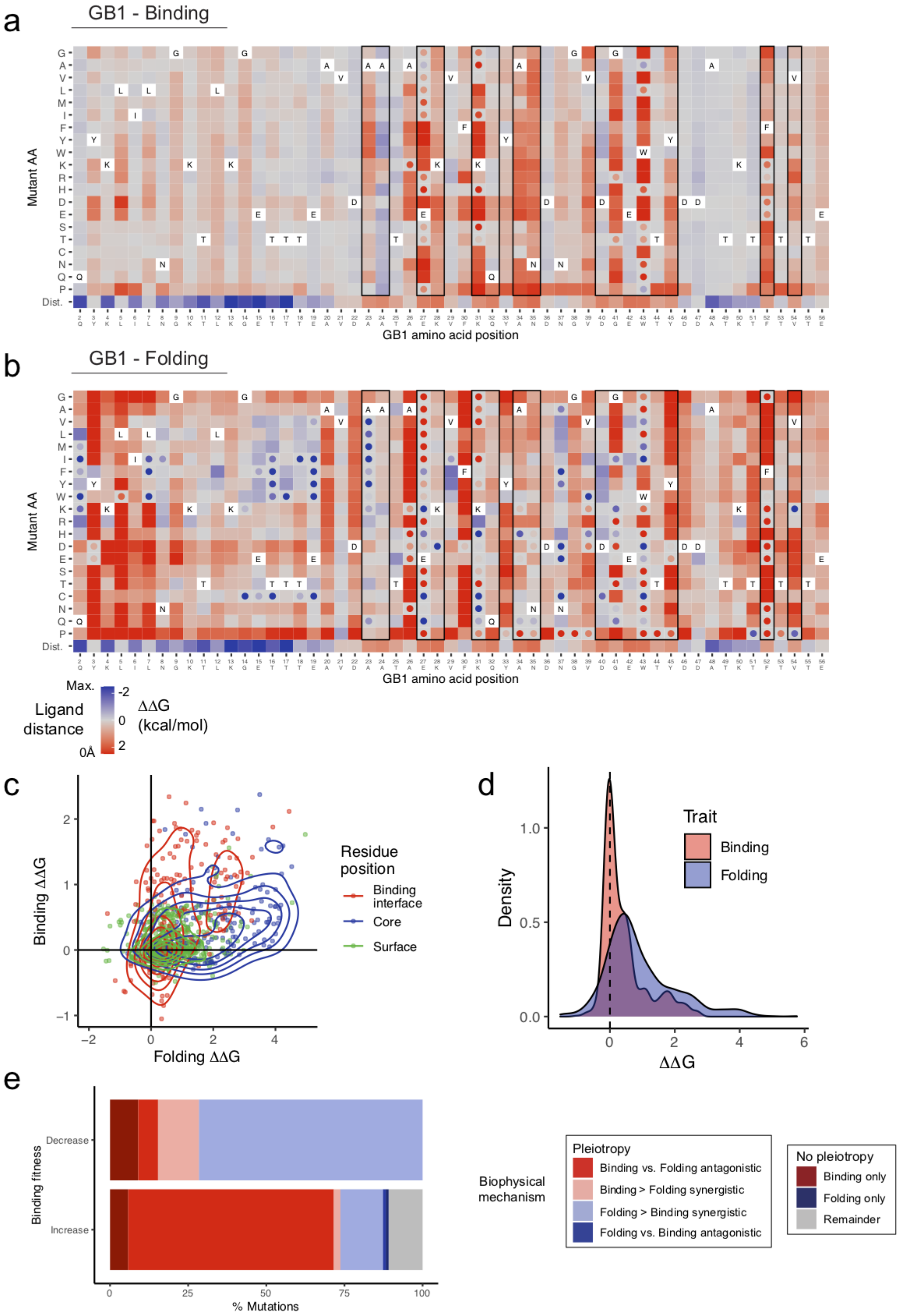

a
GB1 - Binding
Mutant AA
GB1 amino acid position
b
GB1 - Folding
Mutant AA
GB1 amino acid position
Dist.
Max.
Ligand distance
0Å
ΔΔG (kcal/mol)
c
Binding ΔΔG
Folding ΔΔG
Residue position
Binding interface
Core
Surface
d
Density
ΔΔG
Trait
Binding
Folding
e
Binding fitness
Decrease
Increase
% Mutations
Biophysical mechanism
Pleiotropy
Binding vs. Folding antagonistic
Binding > Folding synergistic
Folding > Binding synergistic
Folding vs. Binding antagonistic
No pleiotropy
Binding only
Folding only
Remainder

**Figure S7. Binding and folding free energy landscapes of the GB1 domain.**

**a-b.** Heatmaps showing inferred changes in free energies of binding (**a**) and folding (**b**) for the GB1 domain. The final row in each heatmap indicates the minimal distance to the ligand (considering the side chain heavy atoms or the alpha carbon atoms in the case of glycine). Free energy changes of ligand-proximal residues (ligand distance<5Å) are boxed. Low confidence estimates are indicated with dots (95% confidence interval ≥1kcal/mol). Free energy changes more extreme than ±2.5 were set to this limit. **c.** Scatterplots comparing binding and folding free energy changes of mutations in the core, surface and binding interface. Contours indicate estimates of 2D densities with 6 contour bins. **d.** Distribution of binding (red) and folding (blue) free energy changes. **e.** Percentage of mutations that significantly decrease (top) or increase (bottom) fitness in the binding assay (FDR<0.05) categorized by their biophysical mechanism. Pleiotropic mutations have significant changes in free energies of both folding and binding (FDR<0.05) and are classified as either synergistic or antagonistic depending on whether their effects are in the same or different direction respectively.

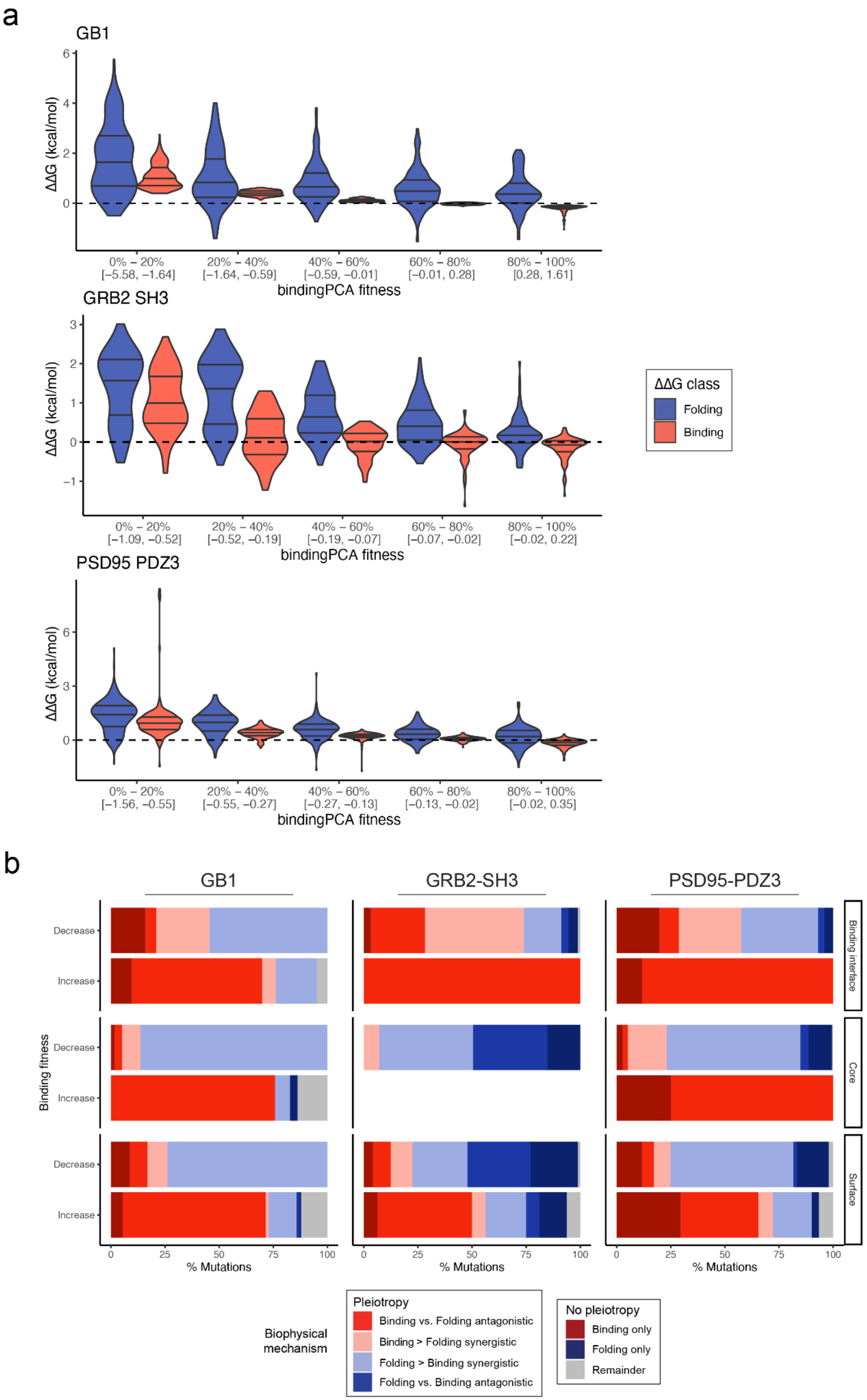
a
GB1
ΔΔG (kcal/mol)
0% – 20% [−5.58, −1.64]
20% – 40% [−1.64, −0.59]
40% – 60% [−0.59, −0.01]
60% – 80% [−0.01, 0.28]
80% – 100% [0.28, 1.61]
bindingPCA fitness
GRB2 SH3
ΔΔG (kcal/mol)
0% – 20% [−1.09, −0.52]
20% – 40% [−0.52, −0.19]
40% – 60% [−0.19, −0.07]
60% – 80% [−0.07, −0.02]
80% – 100% [−0.02, 0.22]
bindingPCA fitness
ΔΔG class
Folding
Binding
PSD95 PDZ3
ΔΔG (kcal/mol)
0% – 20% [−1.56, −0.55]
20% – 40% [−0.55, −0.27]
40% – 60% [−0.27, −0.13]
60% – 80% [−0.13, −0.02]
80% – 100% [−0.02, 0.35]
bindingPCA fitness
b
GB1
GRB2-SH3
PSD95-PDZ3
Binding fitness
Decrease
Increase
Binding interface
Core
Surface
% Mutations
Pleiotropy
Binding vs. Folding antagonistic
Binding > Folding synergistic
Folding > Binding synergistic
Folding vs. Binding antagonistic
No pleiotropy
Binding only
Folding only
Remainder
Biophysical mechanism

**Figure S8. Biophysical mechanism of mutations that affect binding.**

**a.** Changes in free energy of binding (blue) or folding (red) of single AA substitutions with different fitness effects in the binding assay for the three protein domains. **b.** Percentage of core, surface or ligand binding mutations that significantly decrease (top) or increase (bottom) fitness in the binding assay (FDR<0.05) categorized by their biophysical mechanism. Pleiotropic mutations have significant changes in free energies of both folding and binding (FDR<0.05) and are classified as either synergistic or antagonistic depending on whether their effects are in the same or different direction respectively.

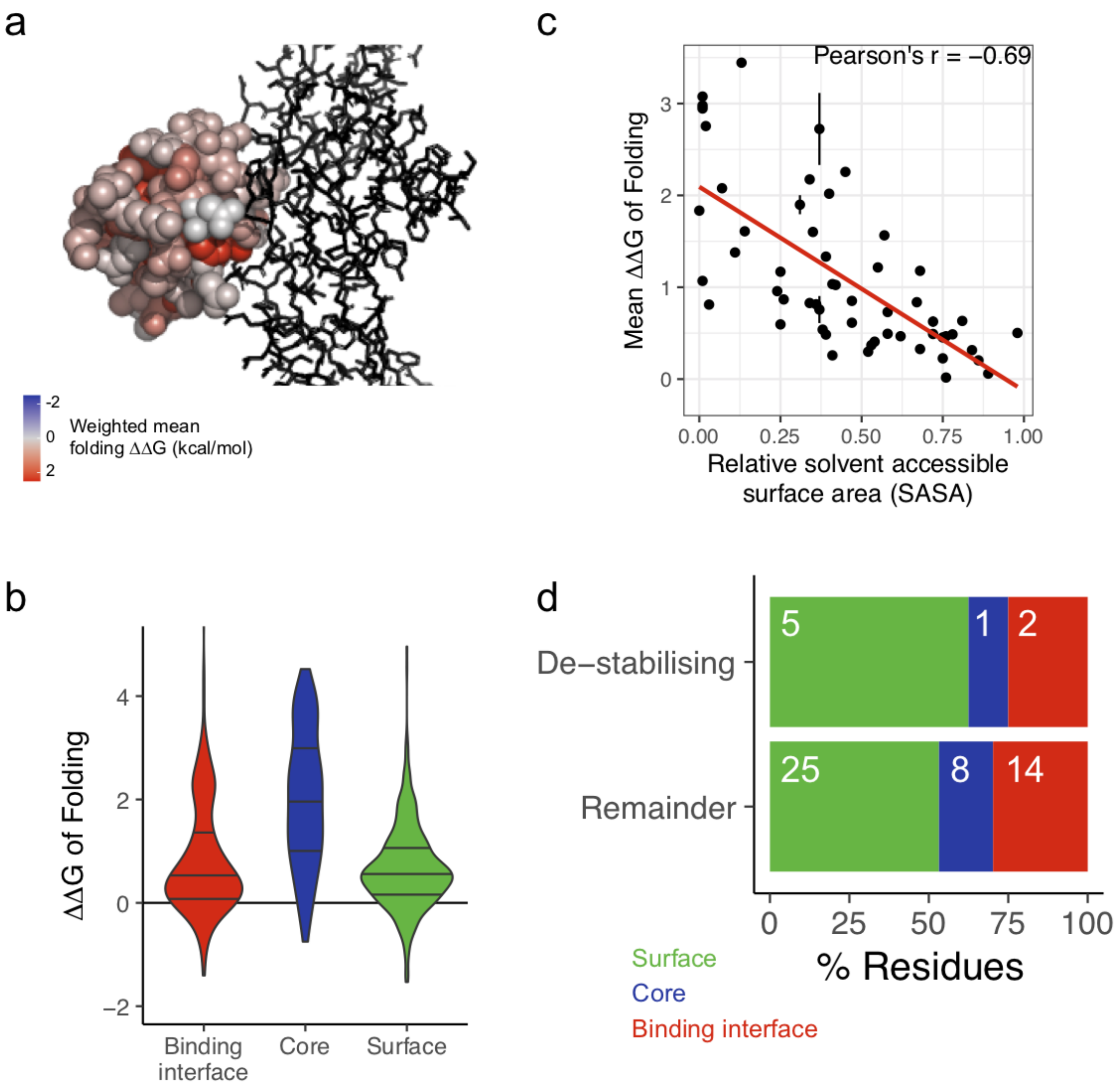


**Figure S9. GB1 mutational effects on protein stability.**

**a**. 3D structure of GB1 (PDB entry 1FCC) where residue atoms are colored by the position-wise average change in the free energy of folding. The FC domain of the human Immunoglobulin G is shown as black sticks. **b**. Violin plots indicating distributions of changes in free energy of folding stratified by position in the structure (two-sided Mann-Whitney U test p-value<2.2e-16 comparing mutations in the core versus the remainder). **c**. Negative correlation between the position-wise average change in free energy of folding and the solvent exposure of the corresponding residue (RSASA). **d**. Percentage of core, surface or binding interface residues shown separately for de-stabilising residues (positions with ≥5 stabilizing mutations, folding ΔΔG<0, FDR<0.05) and the remainder. Inset numbers are total counts.

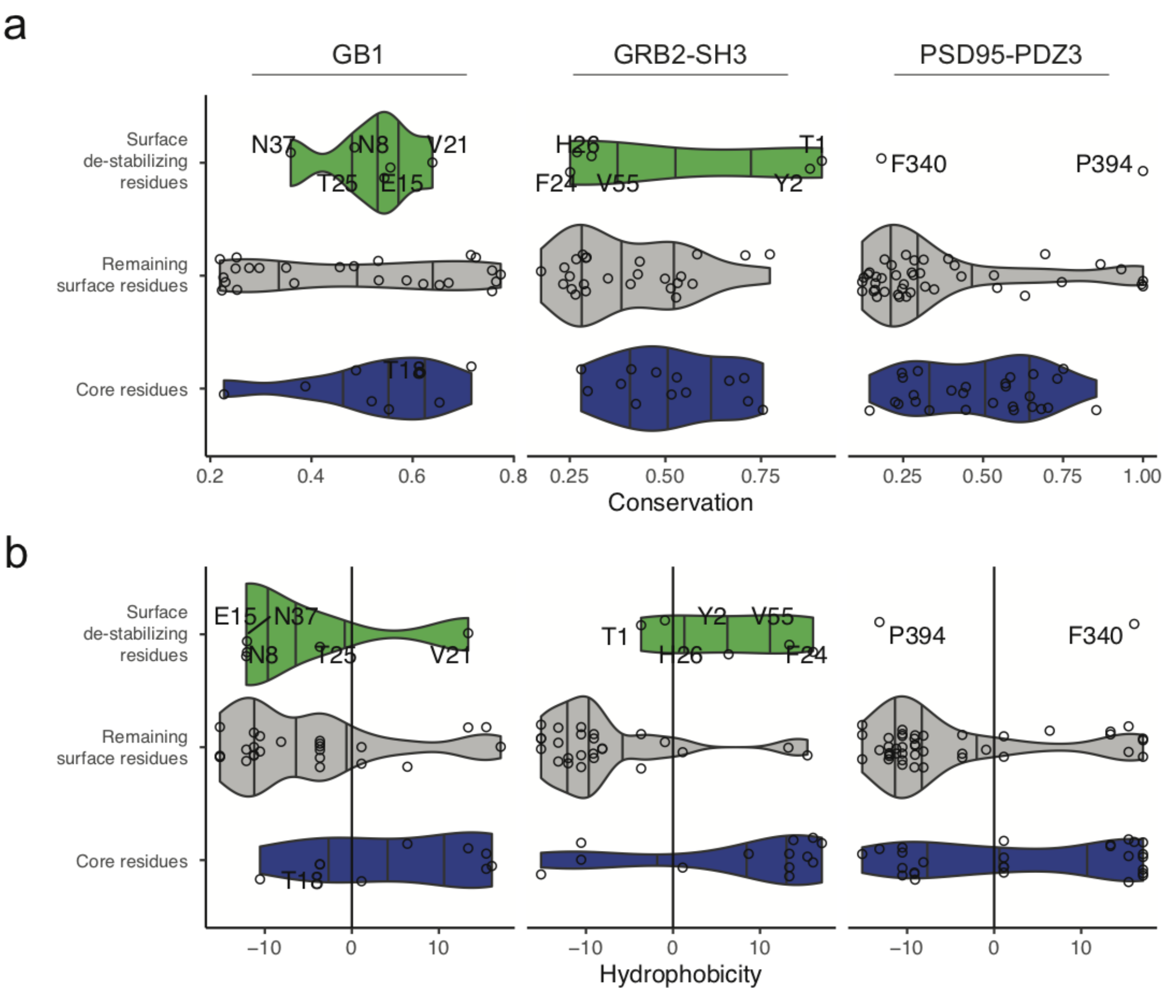


**Figure S10. Evolutionary conservation and hydrophobicity of surface de-stabilising residues.**

**a.** Violin plots indicating evolutionary conservation scores (from a multiple sequence alignment of 185, 8,852, 276,481 homologous sequences of the GB1, GRB2-SH3 and PSD95-PDZ3 domains, respectively) shown separately for surface de-stabilizing residues and remaining surface or core residues. **b.** Violin plots indicating hydrophobicity score distributions (based on principal components analysis of AA properties, see Methods) shown separately for surface de-stabilizing residues and remaining surface or core residues (two-sided Mann-Whitney U test p-value = 3e-3 comparing surface destabilising residues versus the remainder combining residues from the GRB2-SH3 and PSD95-PDZ3 domains).

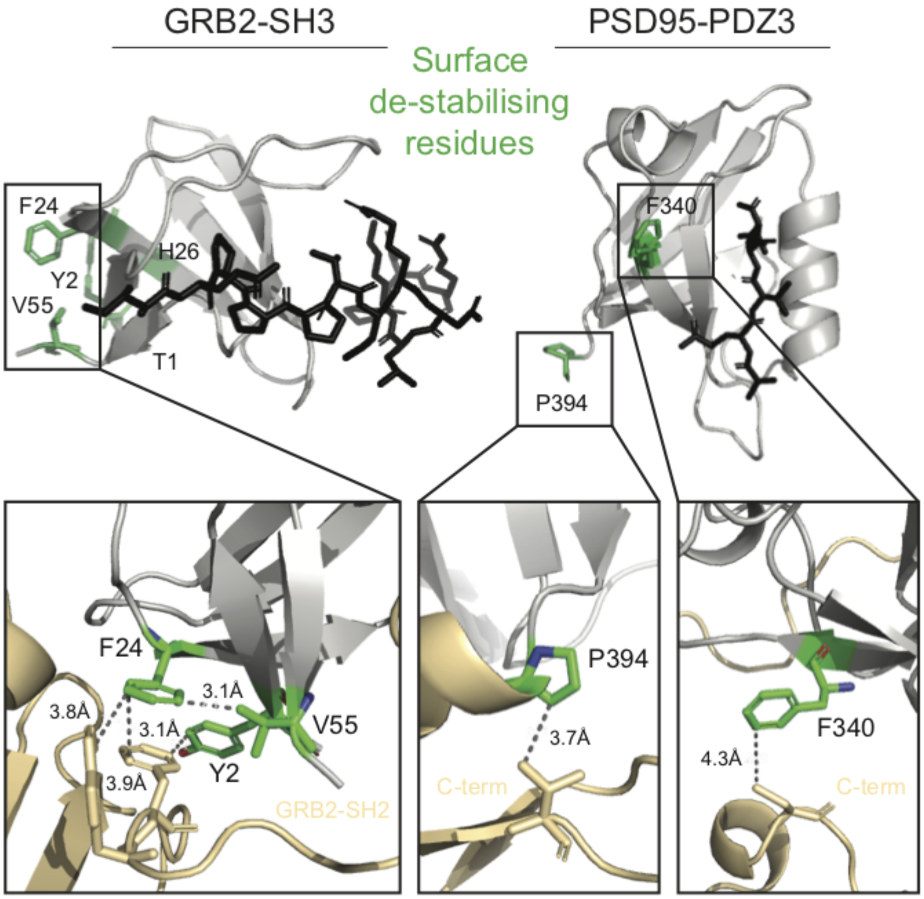


**Figure S11. Surface de-stabilising residues are involved in extra-domain interactions.**

3D structures of the GRB2-SH3 and PSD95-PDZ3 domains (grey cartoons) with the side-chains of surface de-stabilising residues highlighted in green sticks. Ligands are shown as black sticks. In the insets, in yellow is shown the SH2 domains of the second monomer of GRB2 when found in dimeric form (left, PDB entry 1GRI), and relevant proximal portions of PSD95 C-terminal to the PDZ3 domain (middle and right, PDB entry 1BE9 and AlphaFold Protein Structure Database entry P78352).

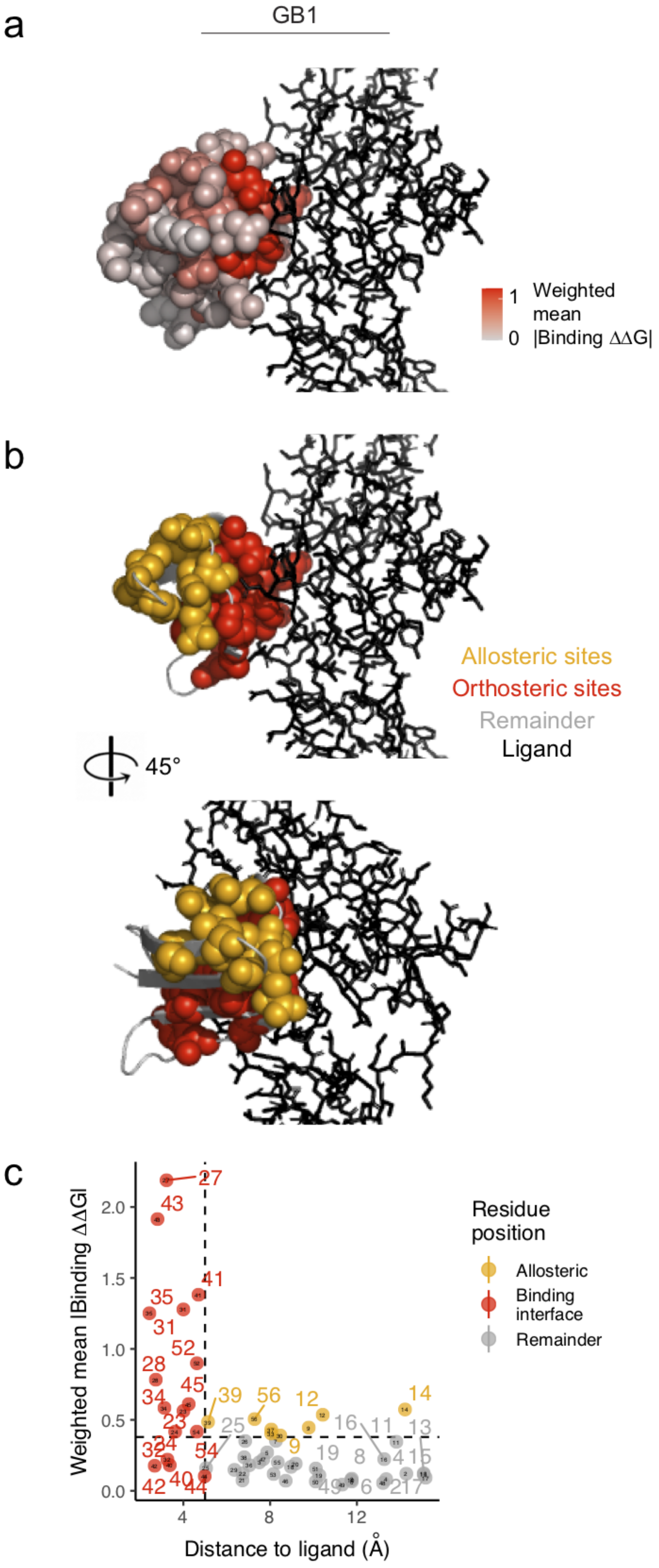
a
GB1
1
0
Weighted mean |Binding ΔΔG|
b
Allosteric sites
Orthosteric sites
Remainder
Ligand
45°
c
Weighted mean |Binding ΔΔG|
Distance to ligand (Å)
Residue position
Allosteric
Binding interface
Remainder

**Figure S12. Major allosteric sites in the GB1 domain.**

**a.** 3D structures of the protein G B1 domain where residue atoms are colored by the position-wise average absolute change in the free energy of binding. The FC domain of the human Immunoglobulin G is shown as black sticks. **b.** GB1 domain structure with binding interface residues (ligand distance<5Å) highlighted in red spheres and major allosteric site residues highlighted in orange spheres **c.** Relationship between the position-wise average absolute change in free energy of binding and the distance to the ligand (minimal side chain heavy atom distance) in the GB1 domain. Major allosteric sites (yellow) are defined as non-binding interface residues with weighted average absolute change in free energy of binding higher than the average of binding interface residue mutations (red).

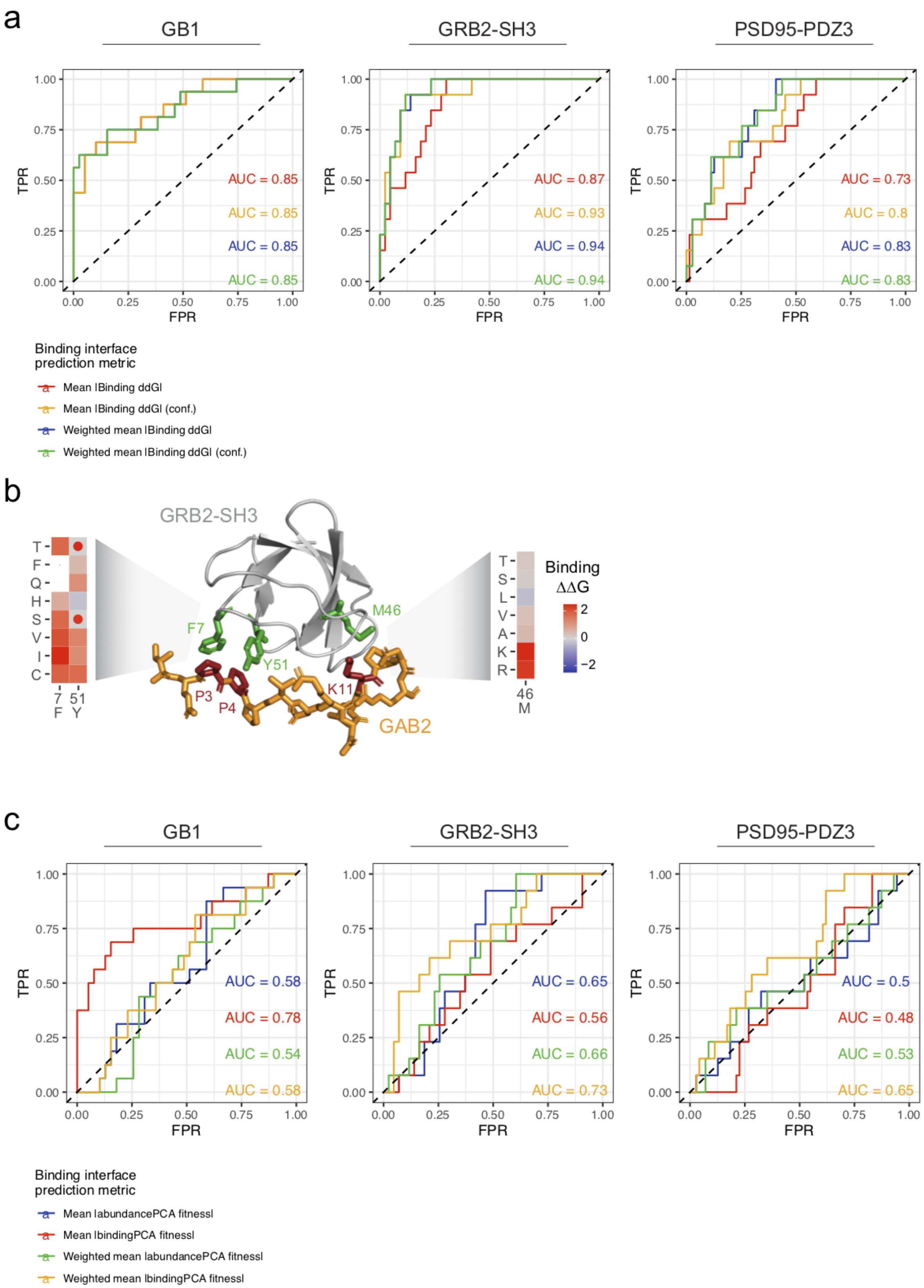
a
GB1
GRB2-SH3
PSD95-PDZ3
TPR
FPR
0.00
0.25
0.50
0.75
1.00
AUC = 0.85
AUC = 0.85
AUC = 0.85
AUC = 0.85
AUC = 0.87
AUC = 0.93
AUC = 0.94
AUC = 0.94
AUC = 0.73
AUC = 0.8
AUC = 0.83
AUC = 0.83
Binding interface
prediction metric
Mean |Binding ddG|
Mean |Binding ddG| (conf.)
Weighted mean |Binding ddG|
Weighted mean |Binding ddG| (conf.)
b
GRB2-SH3
T
F
Q
H
S
V
I
C
7 51
F Y
F7
Y51
M46
P3
P4
K11
GAB2
T
S
L
V
A
K
R
46
M
Binding
ΔΔG
2
0
−2
c
GB1
GRB2-SH3
PSD95-PDZ3
TPR
FPR
0.00
0.25
0.50
0.75
1.00
AUC = 0.58
AUC = 0.78
AUC = 0.54
AUC = 0.58
AUC = 0.65
AUC = 0.56
AUC = 0.66
AUC = 0.73
AUC = 0.5
AUC = 0.48
AUC = 0.53
AUC = 0.65
Binding interface
prediction metric
Mean |abundancePCA fitness|
Mean |bindingPCA fitness|
Weighted mean |abundancePCA fitness|
Weighted mean |bindingPCA fitness|

**Figure S13. Changes in free energy of binding in ligand binding interfaces.**

**a.** ROC curves for predicting ligand contacting residues (<5Å) using (weighted) mean absolute binding ΔΔG considering all variants or those with confident inferred free energies (conf., see Figure S2.4). AUC=Area Under the Curve. **b.** Inferring changes in free energy of binding provides insights into the interactions that mediate binding between GRB2-SH3 and GAB2 peptide, and how mutations disrupt binding. F7 and Y51 of the GRB2-SH3 domain contact P3 and P4 of the GAB2 peptide through aromatic-proline interactions (left heatmap). In these two positions, only mutations to Y, F, Q and H, which can interact with proline through aromatic-proline or amino-aromatic interactions, are tolerated, while all other amino acid substitutions result in decreased binding affinity (positive binding ΔΔG). Residue M46 can tolerate all amino acid substitutions except to positively charged residues (right heatmap). The closest residue of GAB2 is a lysine, and so a repulsive electrostatic interaction likely occurs when a positively charged amino acid occupies position 46 of the SH3 domain (binding ΔΔG of 2.1 and 1.99 for M46K and M46R respectively). **c.** ROC curves for predicting ligand contacting residues using (weighted) mean *bindingPCA* or *abundancePCA* fitness.

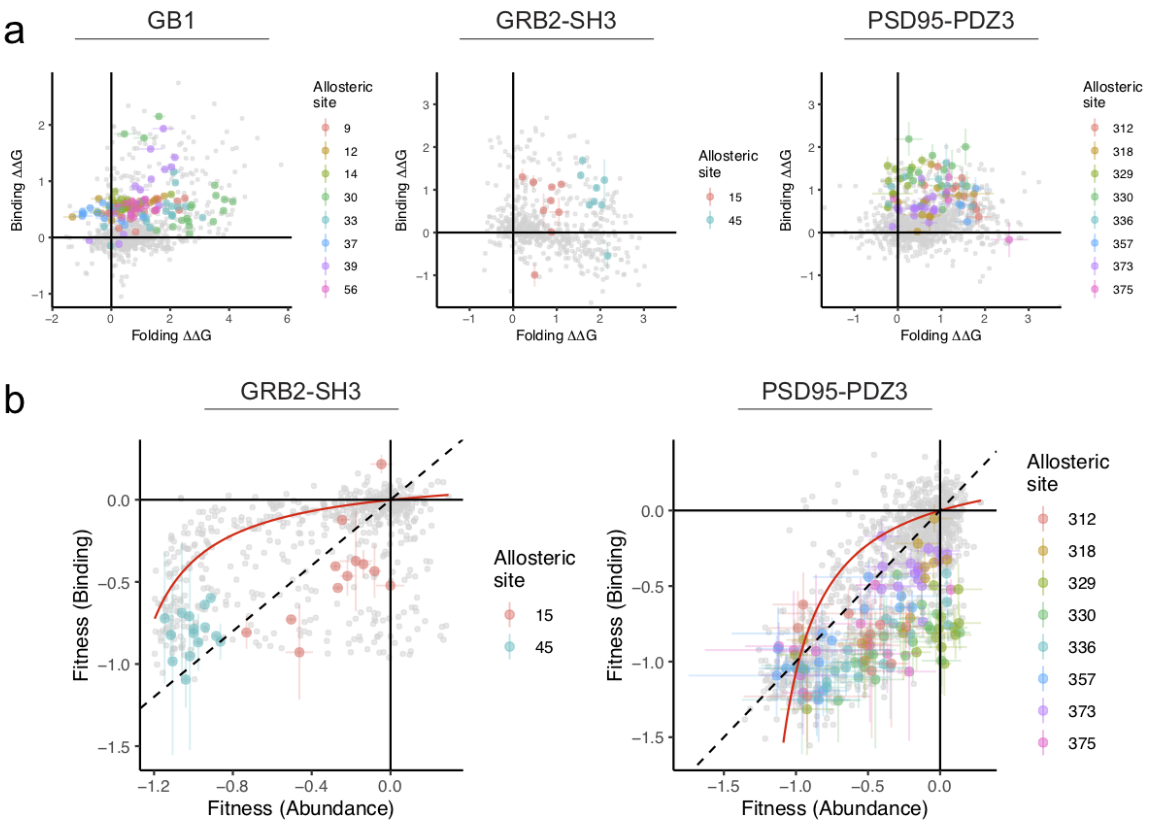


**Figure S14. Changes in fitness and free energy of binding and folding of major allosteric sites.**

**a.** Scatterplots of single AA substitutions' changes in free energy of binding and folding for the GB1 (left panel), GRB2-SH3 (middle panel) and PSD95-PDZ3 (right panel) protein domains. Variants are coloured by AA position if found in a major allosteric site and error bars indicate the 95% confidence range of the inter-fold distribution. **b.** Scatterplots comparing abundance and binding fitness of single AA substitutions in the GRB2-SH3 (left panel) and PSD95-PDZ3 (right panel). Variants are coloured by AA position if found in a major allosteric site. Error bars indicate 95% confidence intervals. The red line indicates the model-derived relationship between abundance and binding fitness in the absence of a change in the free energy of binding.

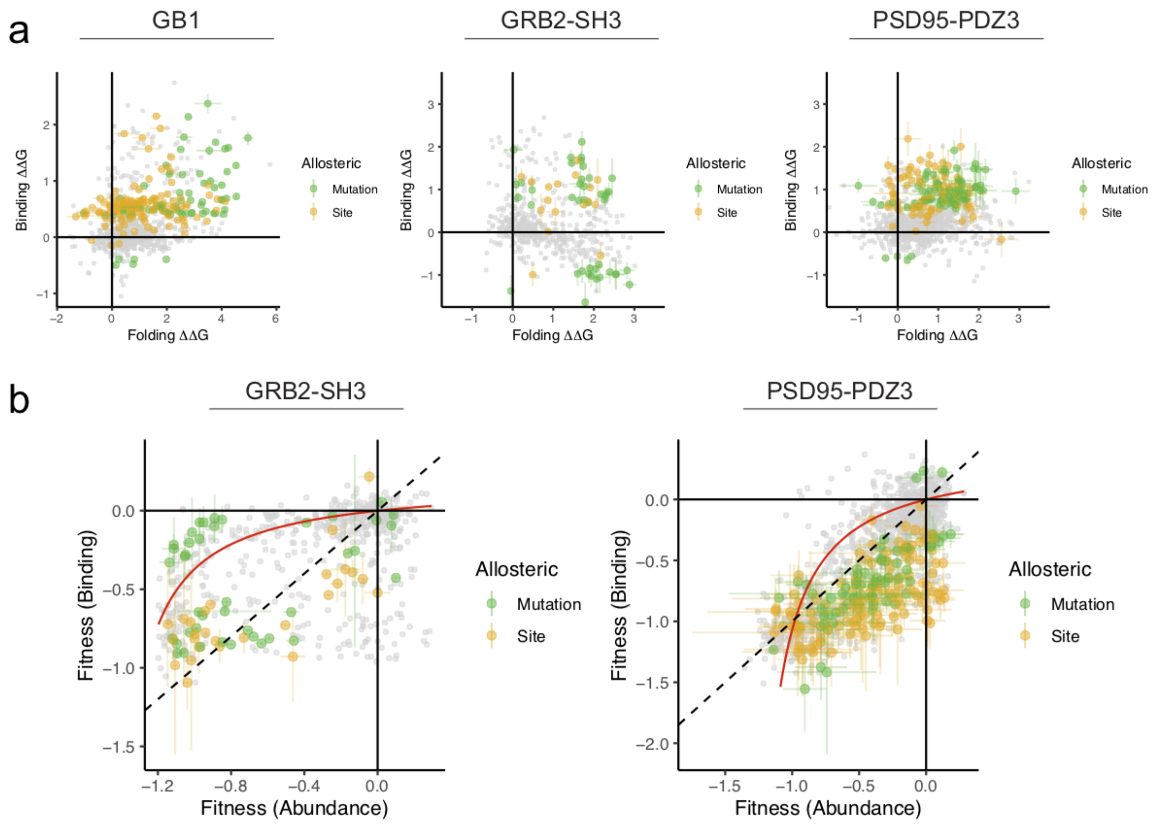


**Figure S15. Changes in fitness and free energy of binding and folding of allosteric sites and positions with allosteric mutations.**

**a.** Scatterplots of single AA substitutions' changes in free energy of binding and folding for the GB1 (left panel), GRB2-SH3 (middle panel) and PSD95-PDZ3 (right panel) protein domains. Variants are coloured by AA position if found in a major allosteric site (yellow) or in a position that has allosteric mutations (green) and error bars indicate the 95% confidence range of the inter-fold distribution. **b.** Scatterplots comparing abundance and binding fitness of single AA substitutions in the GRB2-SH3 (left panel) and PSD95-PDZ3 (right panel). Variants are coloured by AA position if found in a major allosteric site (yellow) or in a position that has allosteric mutations (green). Error bars indicate 95% confidence intervals. The red line indicates the model-derived relationship between abundance and binding fitness in the absence of a change in the free energy of binding.

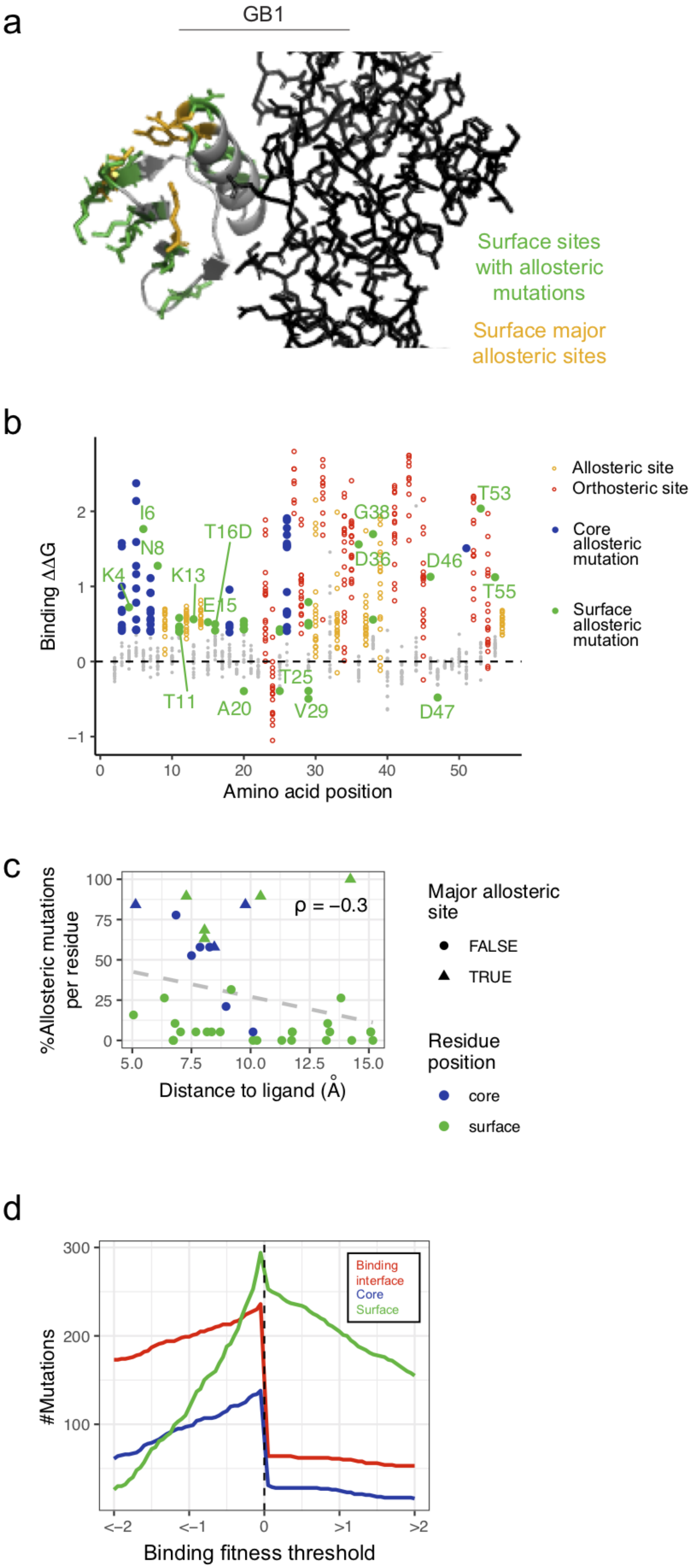
a
GB1
Surface sites with allosteric mutations
Surface major allosteric sites
b
Binding ΔΔG
Amino acid position
K4
I6
N8
T11
K13
E15
T16D
A20
T25
V29
D36
G38
D46
D47
T53
T55
Allosteric site
Orthosteric site
Core allosteric mutation
Surface allosteric mutation
c
%Allosteric mutations per residue
Distance to ligand (Å)
ρ = −0.3
Major allosteric site
FALSE
TRUE
Residue position
core
surface
d
#Mutations
Binding fitness threshold
Binding interface
Core
Surface

**Figure S16. Allosteric mutations in GB1.**

**a**. Domain structure of GB1 with surface allosteric sites and surface residues with allosteric mutations highlighted in orange and green respectively. The FC domain of the human Immunoglobulin G is shown as black sticks. **b**. Scatterplot showing the binding free energy changes of all mutations and coloured according to residue position: allosteric site (orange), orthosteric site/mutation (red), core allosteric mutation (blue), surface allosteric mutation (green). **c.** Percentage of allosteric mutations per residue versus ligand proximity, excluding sites within the binding interface. Points are colored according to residue position and major allosteric sites are indicated (see legend). $\rho$=Spearman rank correlation coefficient. **d.** Total numbers of mutations decreasing or increasing the free energy of binding beyond the indicated minimum or maximum thresholds (x-axis; nominal p-value<0.05) respectively, stratified by position in the structure. Only mutations with confident free energy changes are shown.

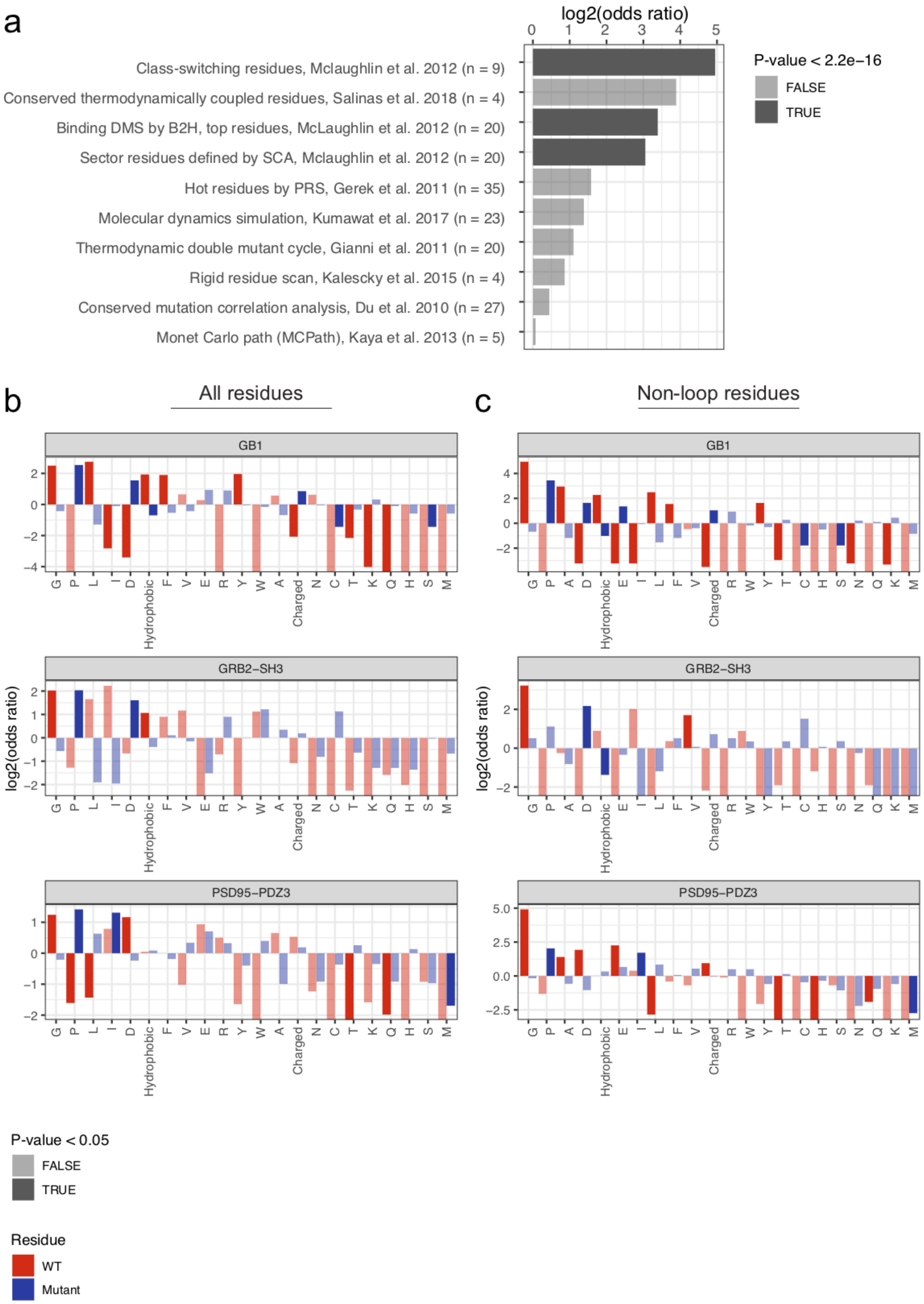
a
log2(odds ratio)
P-value < 2.2e−16
FALSE
TRUE
Class-switching residues, Mclaughlin et al. 2012 (n = 9)
Conserved thermodynamically coupled residues, Salinas et al. 2018 (n = 4)
Binding DMS by B2H, top residues, McLaughlin et al. 2012 (n = 20)
Sector residues defined by SCA, Mclaughlin et al. 2012 (n = 20)
Hot residues by PRS, Gerek et al. 2011 (n = 35)
Molecular dynamics simulation, Kumawat et al. 2017 (n = 23)
Thermodynamic double mutant cycle, Gianni et al. 2011 (n = 20)
Rigid residue scan, Kalescky et al. 2015 (n = 4)
Conserved mutation correlation analysis, Du et al. 2010 (n = 27)
Monet Carlo path (MCPath), Kaya et al. 2013 (n = 5)
b
All residues
c
Non-loop residues
GB1
GRB2−SH3
PSD95−PDZ3
log2(odds ratio)
Hydrophobic
Charged
P-value < 0.05
FALSE
TRUE
Residue
WT
Mutant

**Figure S17. Enrichment of allosteric mutations in literature allosteric networks and specific residue types and classes.**

**a**. Enrichment of allosteric mutations in sets of residues defined by previously reported allosteric networks shown separately for all protein domains. The log2 odds ratio corresponding to a 2x2 contingency table is shown on the x-axis and the associated P-value from a two-sided Fisher's Exact Test is indicated. Residues within the binding interface (ligand distance<5Å) were ignored. Original literature allosteric network sizes are shown in parentheses. **b-c**. Same as (**a**) except sets of residues are defined by the identity of the WT or mutant amino acid (see legend) or their physicochemical properties (hydrophobic i.e. A, V, I, L, M, F, Y, W or charged i.e. R, H, K, D, E). Results are shown for all residues outside the binding interface (**b**) and further restricted to those residues in beta strands or helices i.e. not within loops/turns (**c**). Sets are ranked by their mean effect across the three protein domains.

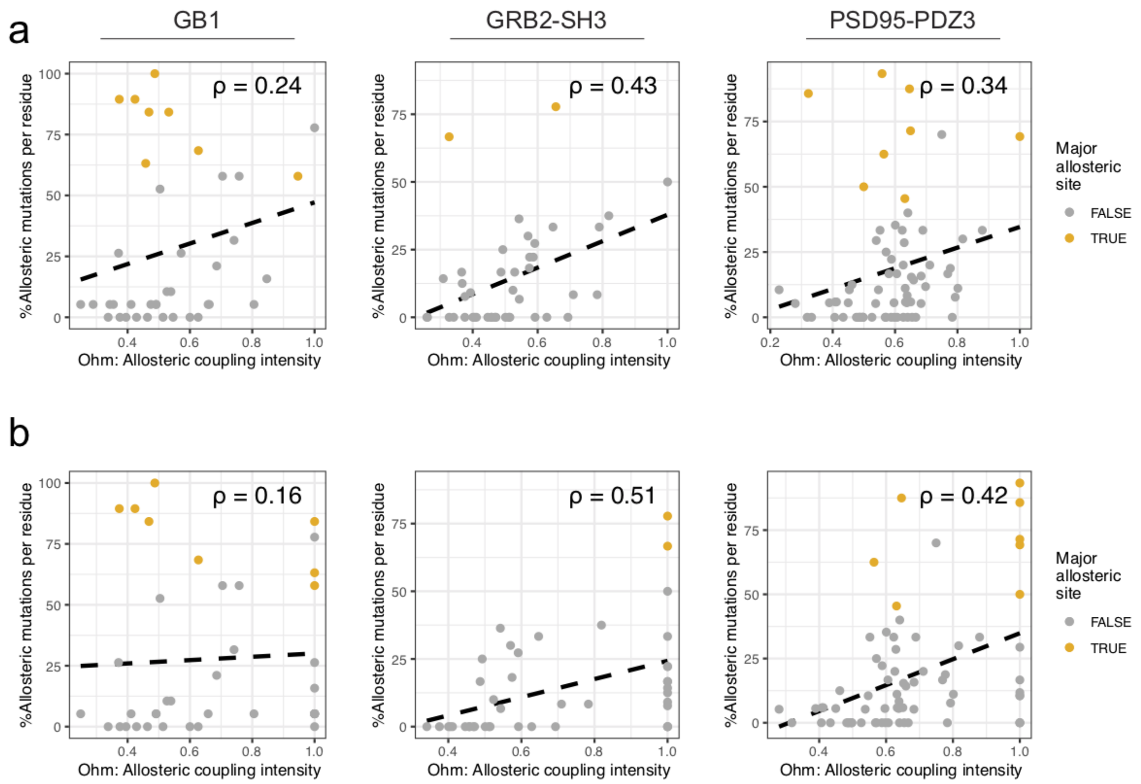


**Figure S18. Comparison of computationally predicted allosteric coupling scores to percentage of allosteric mutations per residue.**

**a**. Percentage of allosteric mutations per residue versus allosteric coupling scores estimated by a network-based perturbation propagation algorithm[7], where residues in the binding interface (ligand distance<5Å) are omitted as they represent the query set. **b**. Same as (**a**) except residues immediately adjacent to binding interface residues in the linear AA sequence (i.e. backbone-backbone contacts which are disregarded by the Ohm algorithm) were given the maximum allosteric coupling score (1.0). Major allosteric sites (in yellow) and Spearman rank correlation coefficients (ρ) are indicated.

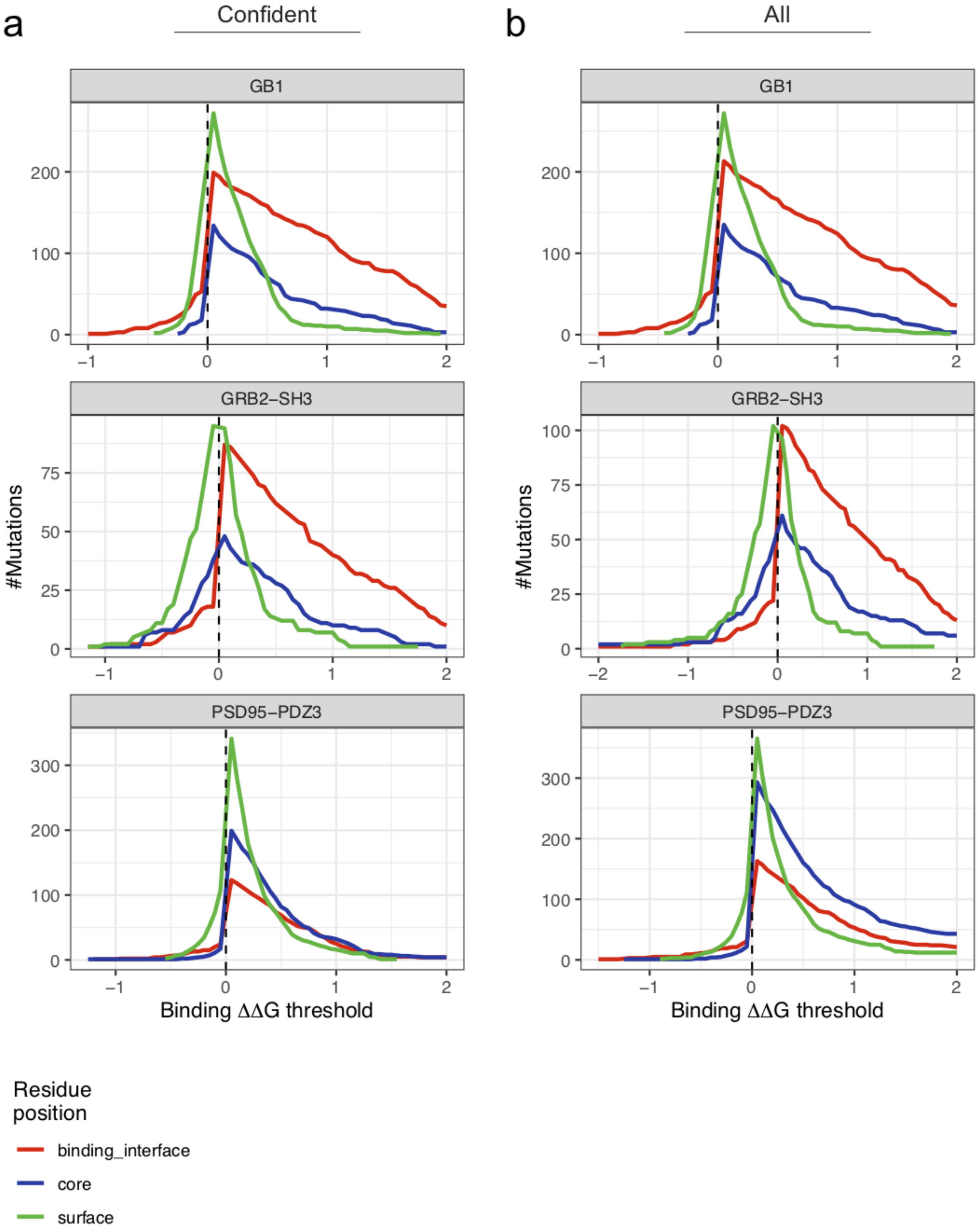
a
Confident
b
All
GB1
GRB2-SH3
PSD95-PDZ3
#Mutations
Binding ΔΔG threshold
Residue position
binding_interface
core
surface

**Figure S19. Mutational biases towards increased or decreased binding given the position in the domain structure.**

**a-b.** Total numbers of mutations decreasing or increasing the free energy of binding beyond the indicated minimum or maximum thresholds (x-axis) respectively, stratified by position in the structure considering (**a**) variants with confident inferred free energies or (**b**) all variants.

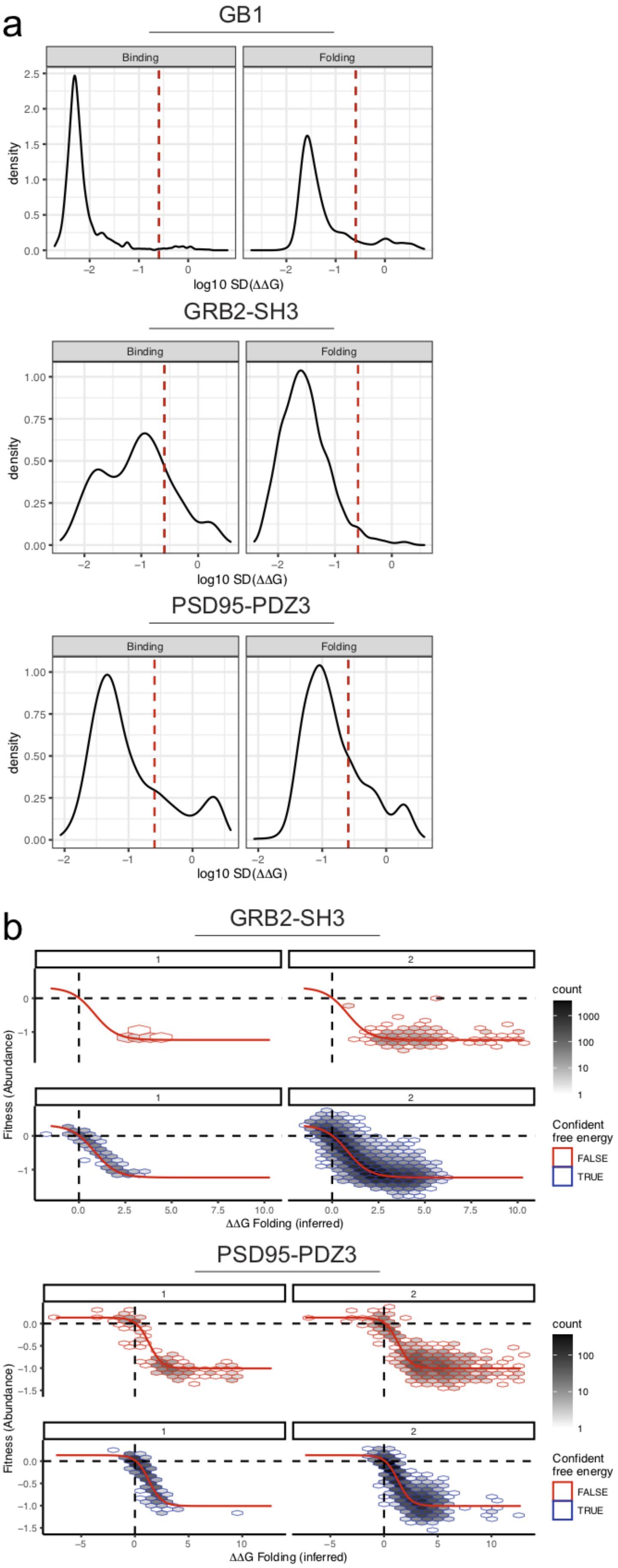
a
GB1
Binding
Folding
density
log10 SD(ΔΔG)
GRB2-SH3
Binding
Folding
density
log10 SD(ΔΔG)
PSD95-PDZ3
Binding
Folding
density
log10 SD(ΔΔG)
b
GRB2-SH3
Fitness (Abundance)
ΔΔG Folding (inferred)
count
Confident free energy
FALSE
TRUE
PSD95-PDZ3
Fitness (Abundance)
ΔΔG Folding (inferred)
count
Confident free energy
FALSE
TRUE

**Figure S20. Determining confident inferred free energy changes using a Monte Carlo simulation approach.**

**a.** Distributions of inferred binding (left) and folding (right) free energy change variability (standard deviation) calculated between ten models fit using data from [1] independent random samples of fitness estimates from their underlying error distributions and [2] independent random samples of the validation data consisting of 10% of double AA substitution variants held out during training. Confident inferred free energy changes are defined as those with Monte Carlo simulation derived 95% confidence intervals <1kcal/mol (red dashed line). **b.** 2d density plots showing the non-linear relationships between observed *abundancePCA* fitness and changes in free energy of folding of high (blue) and low confidence (red) single (left panels) and double mutants (right panels). Model predicted fitness is shown (red line).
